# New observations of fluorescent organisms in the Banda Sea and in the Red Sea

**DOI:** 10.1101/2023.09.22.559034

**Authors:** Lars Henrik Poding, Peter Jägers, Budiono Senen, Gino Valentino Limmon, Stefan Herlitze, Mareike Huhn

## Abstract

Fluorescence is a widespread phenomenon found in animals, bacteria, fungi, and plants. In marine environments fluorescence has been proposed to play a role in physiological and behavioral responses. Many fluorescent proteins and other molecules have been described in jellyfish, corals, and fish. Here we describe fluorescence in marine species, which we observed and photographed during night dives in the Banda Sea, Indonesia, and in the Red Sea, Egypt. Among various phyla we found fluorescence in sponges, molluscs, tunicates, and fish. Our study extends the knowledge on how many different organisms fluoresce in marine environments. We describe the occurrence of fluorescence in 27 species, in which fluorescence has not been described yet in peer-reviewed literature. It especially extends the knowledge beyond Scleractinia, the so far best described taxon regarding diversity in fluorescent proteins.

## Introduction

Fluorescence is the property of a molecule to absorb light of a certain wavelength followed by the emission of light with longer wavelength. In bacteria, plants, fungi, and animals, various fluorescent molecules have been described such as chitin, minerals, carotenoids, flavonoids, porphyrins, chlorophyll, phycobiliproteins or green fluorescent protein (GFP) and (GFP)-like fluorescent proteins [1,2].

In marine environments many fluorescent species have been described and for some of these species an ecological, behavioral, and/or physiological function has been suggested [2,3]. In the jellyfish *Aequorea victoria*, for example, where the first fluorescent protein (avGFP) was characterized, GFP is expressed as an acceptor molecule for bioluminescent light [4]. So far fluorescent proteins (FP) have been found in Cnidaria, Vertebrata, Cephalochordata (lancelets), and Arthropoda, but have not been described (i.e. published in peer-reviewed literature) in Porifera (sponges), Ctenophora (comb jellies), Tunicata (tunicates), Hemichordata (hemichordates), Echinodermata (echinoderms), Mollusca (molluscs), Annelida (segmented worms), and Nematoda (roundworms) [2,5–10]. Most fluorescent proteins with blue to red emission spectra (GFP-like fluorescent proteins) have been identified in Anthozoa (e.g. corals and anemones) [5]. There, the fluorescence has been hypothesized to be involved in attraction of photosynthetic symbionts, in photoprotection of the symbionts, and in converting blue light to longer wavelength light in mesophotic environments (photoenhancement) [2,3] where the green to red components of the sunlight are selectively removed with water depth resulting in a blue light environment [2,3]. In fish, red fluorescent body coloration and red fluorescent iris have been proposed to be involved in intraspecific communication [11–15]. In addition, differences in the fluorescence pattern between males and females have been suggested to play a role in sexual communication [12,16].

In contrast to jellyfish, corals, anemones, copepods, lancelets, eels, and moray eels, where fluorescence is based on the expression of fluorescent proteins, non-FP based fluorescence has been observed in Porifera (sponges), Heterobranchia (slugs and snails), Annelida (segmented worms), Crustacea (crustaceans), Osteichthyes (bony fish), and Chondrichthyes (cartilaginous fishes) [7,17–21].

Here we show photographs of new cases of fluorescence in marine species, which were taken during night dives in the Banda Sea, Banda Islands, Indonesia and the Red Sea, Dahab, Egypt. These fluorescent species include nudibranchs, tunicates, sponges, various other invertebrates, and fish. In addition to these new records of fluorescence in marine species, we show detailed Leica THUNDER microscopy images of known fluorescent species to provide precise information about the distribution of fluorescence in different body parts.

## Methods

### Ethics statement

No permits were required as no specimen were collected and photos were taken during recreational Scuba dives.

### Photo acquisition in the field

Images of fluorescent species were acquired during SCUBA diving in the Red Sea in Dahab, Egypt (28°29’20.0”N 34°30’57.2”E) and in the Banda Sea at the Banda Islands, Maluku, Indonesia (4°30’55.62”S 129°53’35.92”E). All dives were started after sunset and limited to a maximum depth of 15 meters, which was monitored by a dive computer. Field observations were carried out in May 2019 (Red Sea) and in February-March 2019 and September 2022 (Banda Sea).

Fluorescence was excited with “Sola” light source (NightSea, USA; ʎ max = 449 nm), Flashlight (D2000; INON; Japan), Ex-Inon excitation filter (NightSea, USA; ʎ max = 454 nm) (Fig.1) or “Blue Star” light source (NightSea, USA; ʎ max = 470 nm). The plastic transmission filter (BF 2, NightSea, USA) in front of the underwater housing showed a low cut at ʎ < 500 nm (Fig.1). Fluorescence photography was performed with Powershot G15 (Canon, Japan) placed in a WP DC 48 (Canon, Japan) underwater housing or Sony α 6000 (Sony, Japan) placed in a SF-A6500 (Seafrogs, Hongkong) underwater housing (details on emission and transmission spectra of light sources and filter specified in Figure 1, details on camera specifications and settings in supplementary 1). All images were processed with Lightroom (Version 2015.1.1, Adobe, USA) and Corel Draw (Version 20.0.0.633, Corel Corporation, Canada), by correcting the brightness and contrast uniformly throughout the entire image, and cropping the images (example of digital processing provided in supplementary 1).

**Figure 1:**
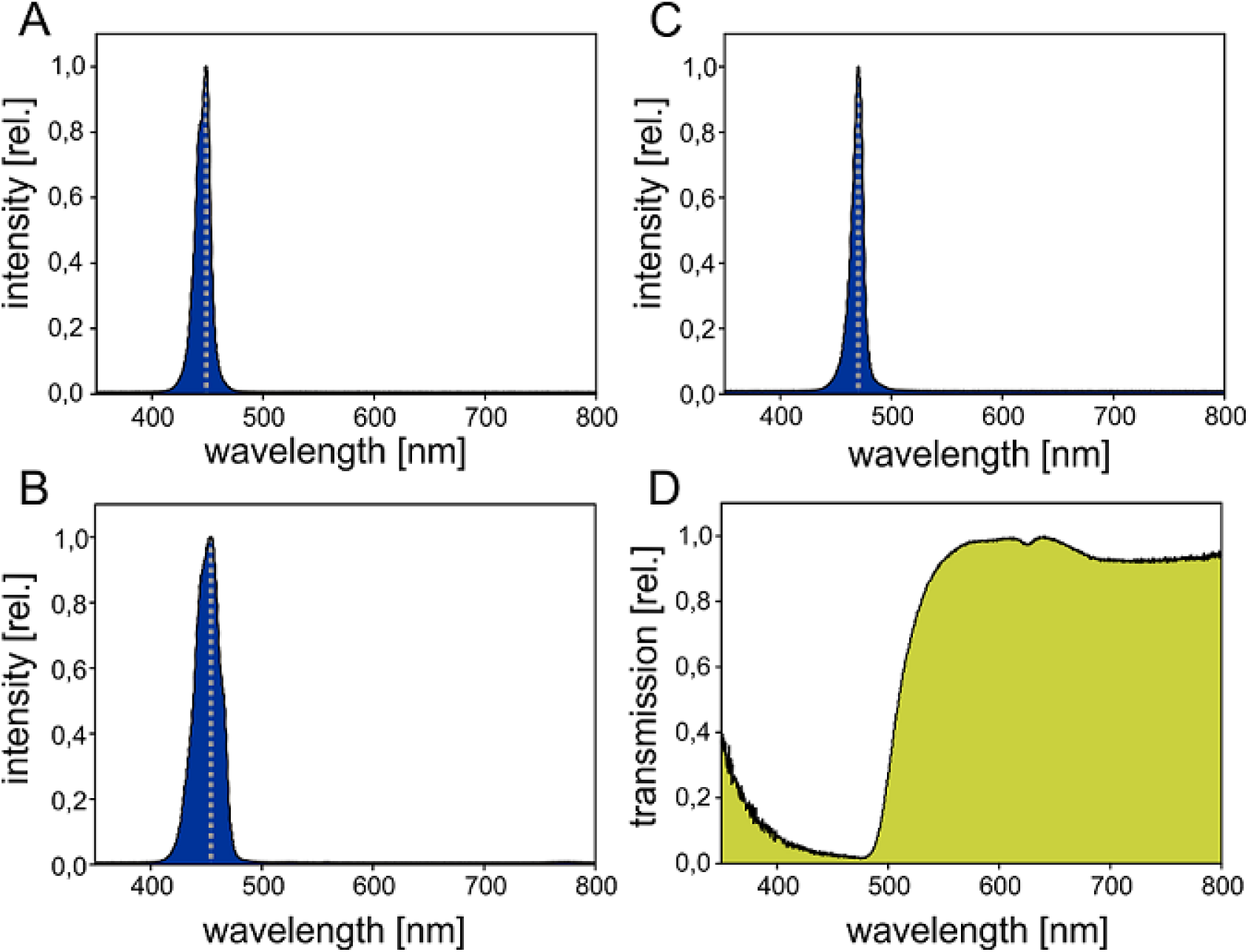
Fluorescence was excited with “Sola” light source (NightSea, USA; ʎ _max_ = 449 nm; A), Flashlight (D2000; INON; Japan) with Ex-Inon excitation filter (NightSea, USA; ʎ _max_ = 454 nm; B) or “Blue Star” light source (NightSea, USA; ʎ _max_ = 470 nm; C). Transmission of the yellow plastic filter BF 2 (NightSea, USA; E) showing a low cut at ʎ < 500 nm. Relative intensity (A-C) and transmission (D) were determined with a Spectrometer (Flame S-UV-VIS-ES, Ocean Optics, USA). A halogen Light Source (HL-3 plus, Ocean Optics, USA) was used to measure transmission of yellow long pass filter.

Species were identified with standard literature [22–31]. The species were identified by comparison of digital underwater pictures with the description and pictures in the literature.

### Photo acquisition by Leica THUNDER microscopy

Detailed images of *Anampses meleagrides, Cirrhilabrus aquamarines, Cirrhilabrus rubrisquamis, Doryrhamphus excisus*, *Eviota atriventris*, *Eviota nigriventris, Lybia tessellata, Nembrotha kubaryana, Odontodactylus scyllarus, Synchiropus sycorax*, and one species of Polyplacophora were acquired with the fluorescence THUNDER stereo microscope Leica M205 FCA equipped with a DFC90000 GT Camera and a GFP (ʎ _Ex_: 395– 455 nm, ʎ _Em_: 480 long pass filter), mCherry (ʎ _Ex_: 540-580 nm, ʎ _Em_: 593-667 nm), and a CY5 (ʎ _Ex_: 590–650 nm, ʎ _Em_: 663–737 nm) filter cube. The fish and invertebrates were purchased from the wholesale trader DeJong Marinelife (Netherlands) in 2021/2022 or from Korallenfarm Joe & Co (Germany). All images were processed with LASX (Version 3.7.6.25997, Leica, Germay), Lightroom (Version 2015.1.1, Adobe, USA), and Corel Draw (Version 20.0.0.633, Corel Corporation, Canada) by correcting the brightness and contrast uniformly, and cropping the images.

## Results

We could identify fluorescence in 27 marine species that had not been described to be fluorescent so far. The species in which the new fluorescent signals were observed belonged to the phyla Porifera, Mollusca, Arthropoda, Annelida, and Chordata.

Porifera – We found green fluorescent spots on the surface of the sponge *Gelliodes fibulata* (Fig. 2 A, B). Furthermore, we observed yellow and orange fluorescence in different unidentified sponges in the Red Sea (Fig. 2 C) and Banda Sea (Fig. 2 D).

**Figure 2:**
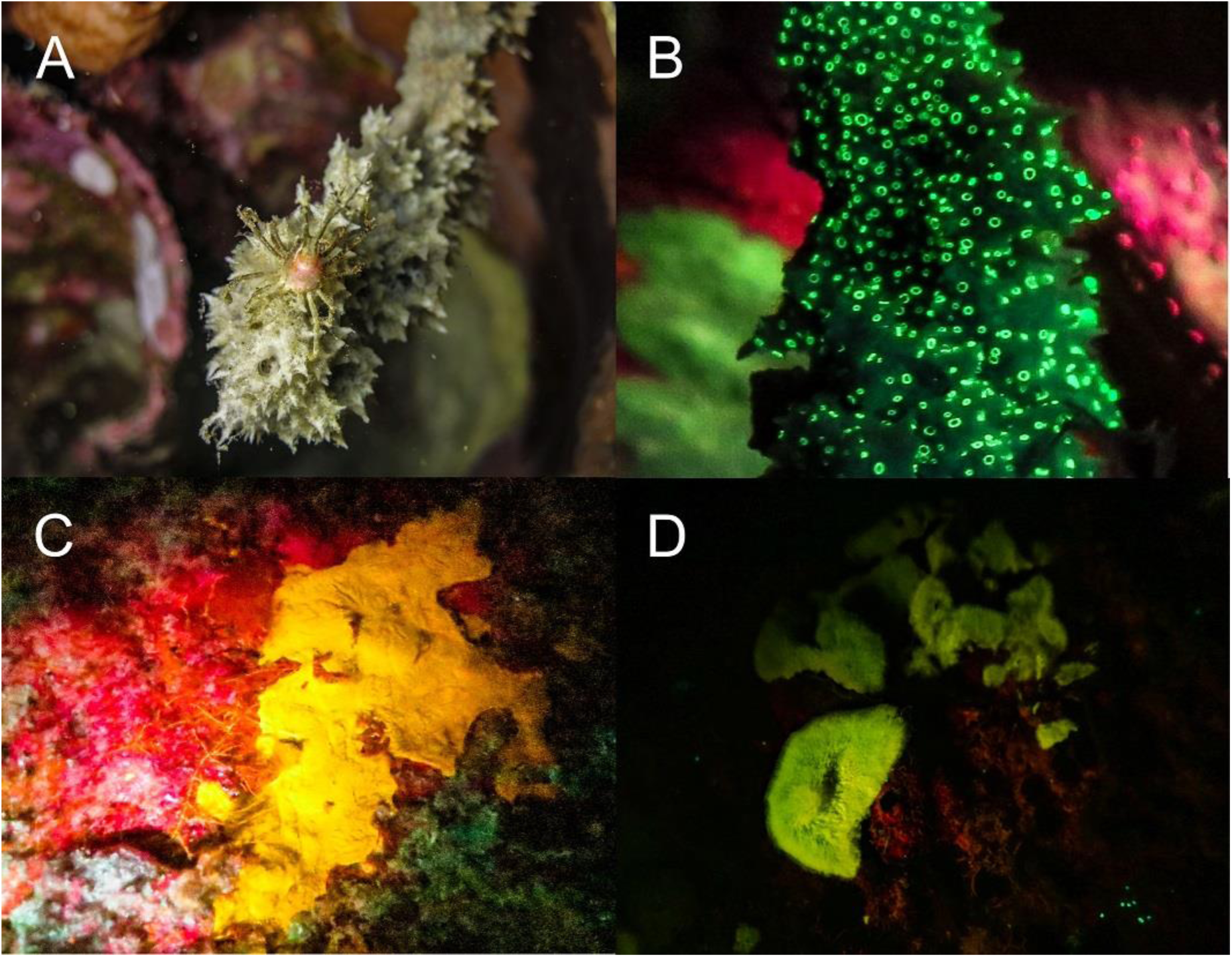
Fluorescence in Porifera in the Banda Sea: The thorny stem sponge *Gelliodes fibulata* ((A & B) white light in A, fluorescence in B) shows green fluorescent spots. Two unidentified sponges (C, D) fluoresce yellow and green.

Scleractinia – We found six species of stony corals that exhibited yellow fluorescence (Fig. 3), *Physogyra lichtensteini,* three different species of *Montipora,* and two *Cycloseris/Fungia* species.

**Figure 3:**
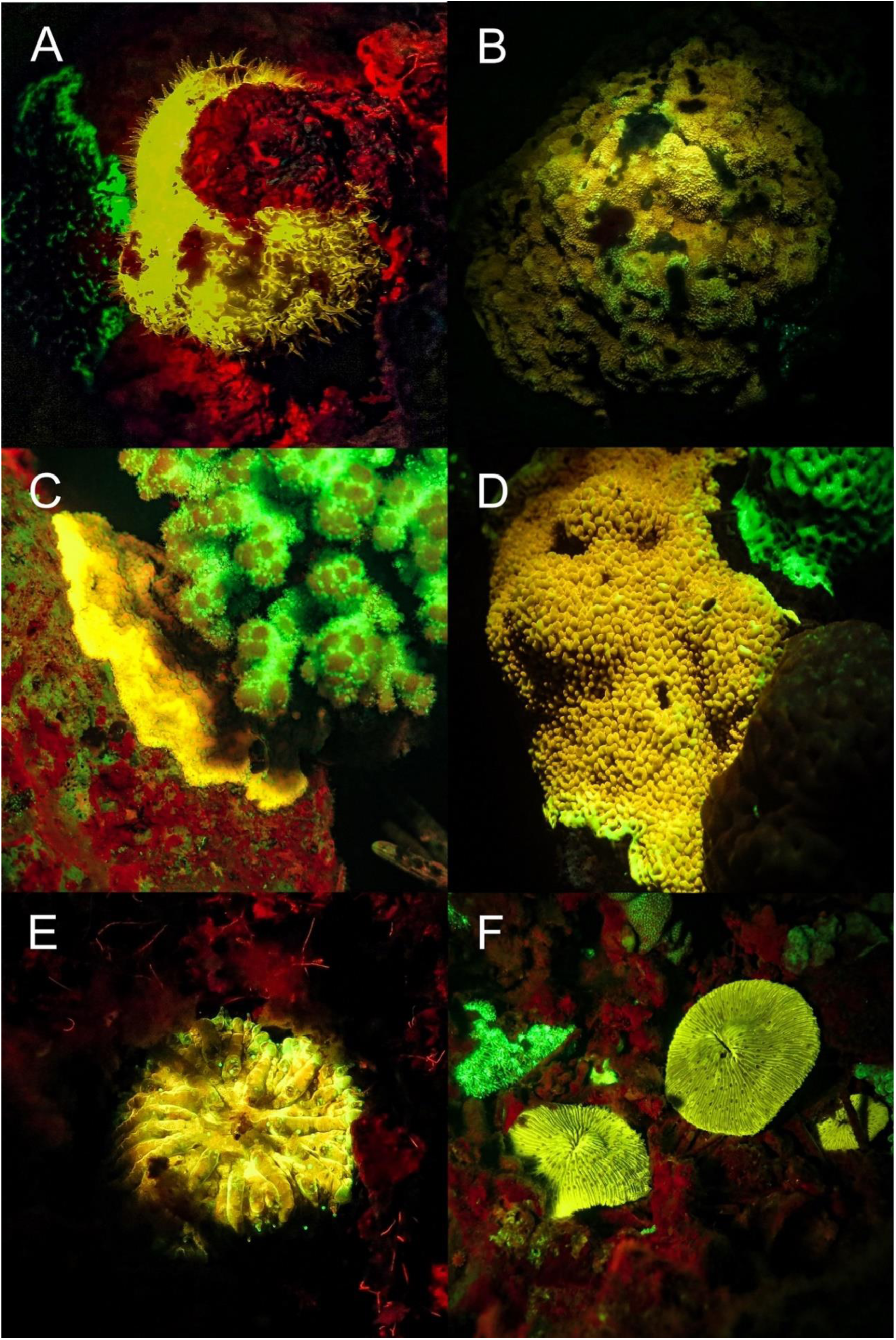
Fluorescence in corals: Yellow fluorescent corals in the Banda Sea and in the Red Sea. *Physogyra lichensteinii* (A), *Montipora sp.* (B - D), and *Cycloseris* sp./*Fungia sp.* (E & F). A, B, D – F: Banda Sea, C: Red Sea.

Molluscs - In molluscs we found fluorescence in the families Octopodidae, Strombidae, Cardiidae, Facelinidae (Fig. 4), Polyceridae (Fig. 5), and unidentified families of Polyplacophora (Fig. 6). The octopus *Abdopus aculeatus* revealed orange fluorescent markings on its body. Bright fluorescent patches were located below the yellow fluorescent eyes and on every arm (Fig. 4 A & B). Other species (unspecified) of the same family showed no fluorescent body markings, but had yellow fluorescent eyes (Fig. 4 B, D, E). We found green and red fluorescence in several snails. The bubble conch *Euprotomus bulla* had a bright green fluorescent operculum and weak green fluorescent eyes. The giant clam *Tridacna crocea* showed bright red fluorescence. (Fig. 4 G). In addition, the nudibranch *Facelina rhodopos* (Fig. 4 H) showed bright orange fluorescence at the tip of the cerata. Another nudibranch *Nembrotha kubaryana* that we photographed in the Banda Sea in its natural habitat (Fig. 5 A) and had purchased from DeJong Marinelife (Netherlands) to image with THUNDER microscopy (Fig. 5 B) revealed strong red fluorescence. We were able to localize the fluorescence in more detail under laboratory settings and found fluorescent patches around the mouth (Fig. 5 D), the rhinophores (Fig. 5 E - G), gills (Fig. 5 B), and the dorsal rim (Fig. 5 B & H). We also found a variety of different Polyplacophora that exhibited bright green, yellow, and red fluorescence (Fig. 6 A - D).

**Figure 4:**
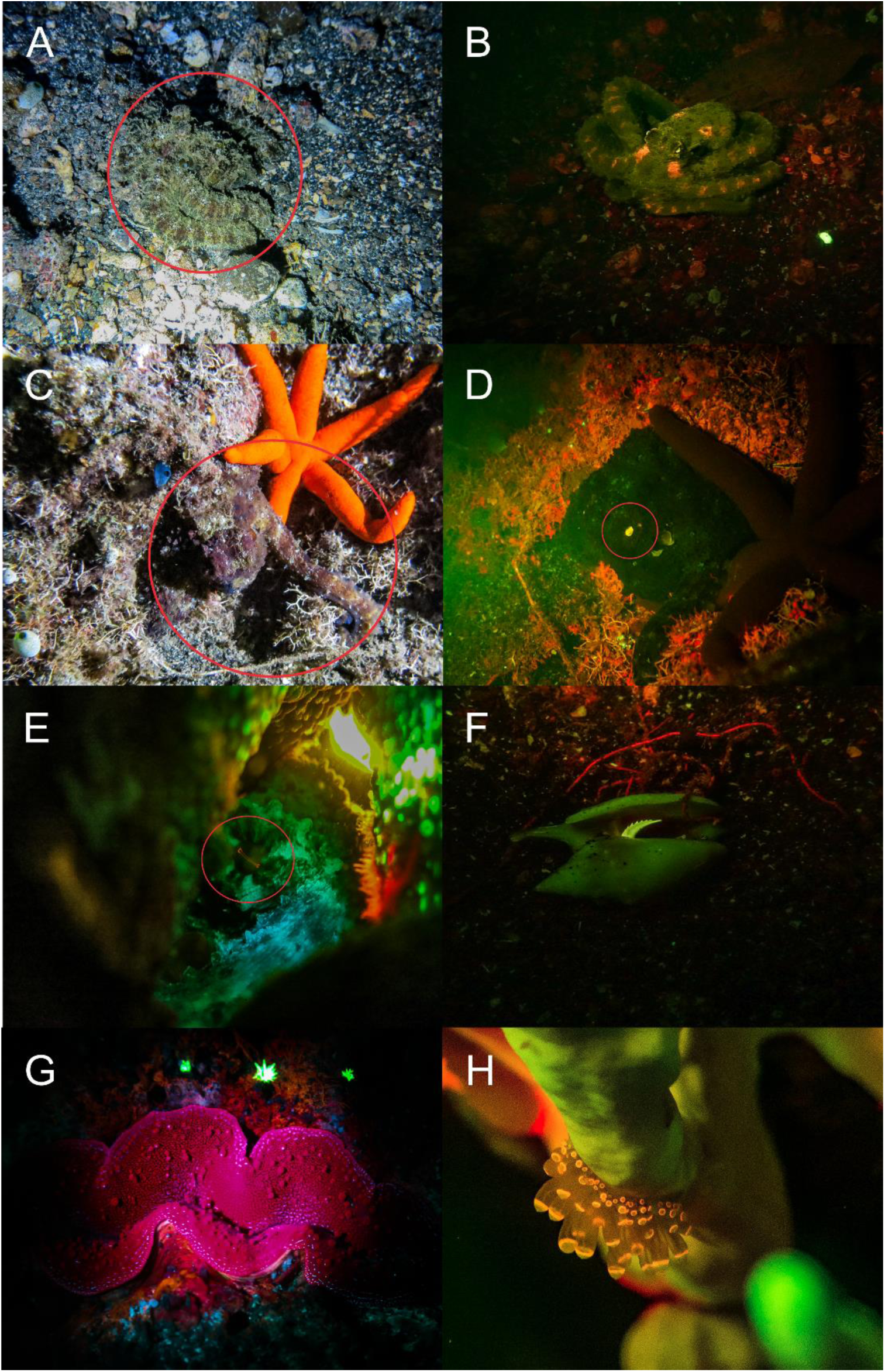
Fluorescence in Mollusca: Three different types of fluorescent Octopods; *Abdopus aculeatus* (A (white light) & B (fluorescence)) with orange fluorescent skin patches, two unidentified species of Octopodidae (C (white light) – E (fluorescence)) with fluorescent eyes, green fluorescent operculum of *Euprotomus bulla* (F), red fluorescent mantle of *Tridacna crocea* (G), orange fluorescent *Facelina rhodopus* (H) on *Millepora sp*.. A – G: Banda Sea, H: Red Sea

**Figure 5:**
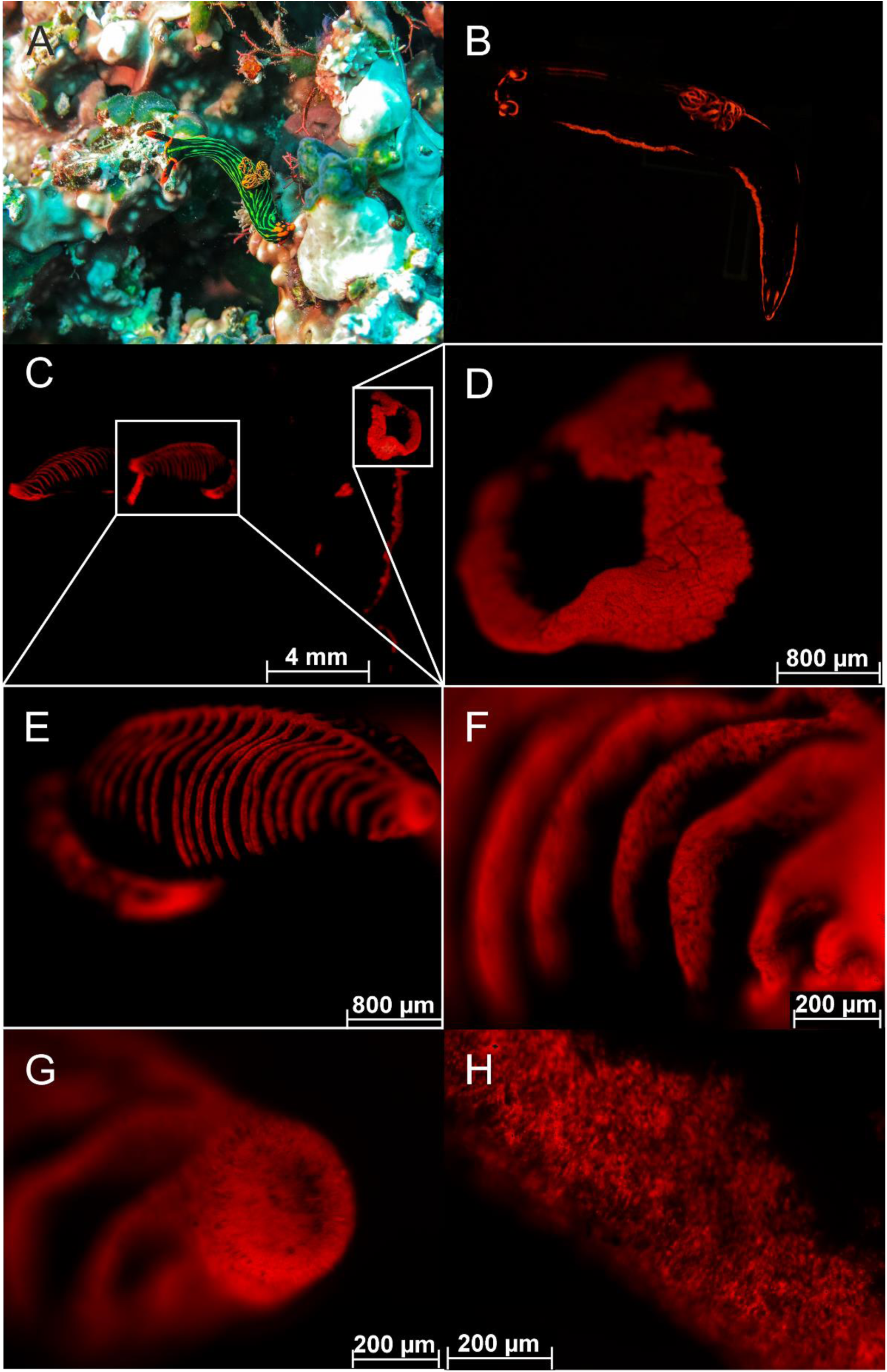
Fluorescence in the nudibranch *Nembrotha kubaryana:* (A) natural habitat (Banda Sea), white light & Fluorescence (B), whole body (B, C), oral tentacles (D), rhinophores (E-G), gill (H) fluorescence. Pictures C – H created with a THUNDER microscope (Leica, Germany) with CY5 filter.

**Figure 6:**
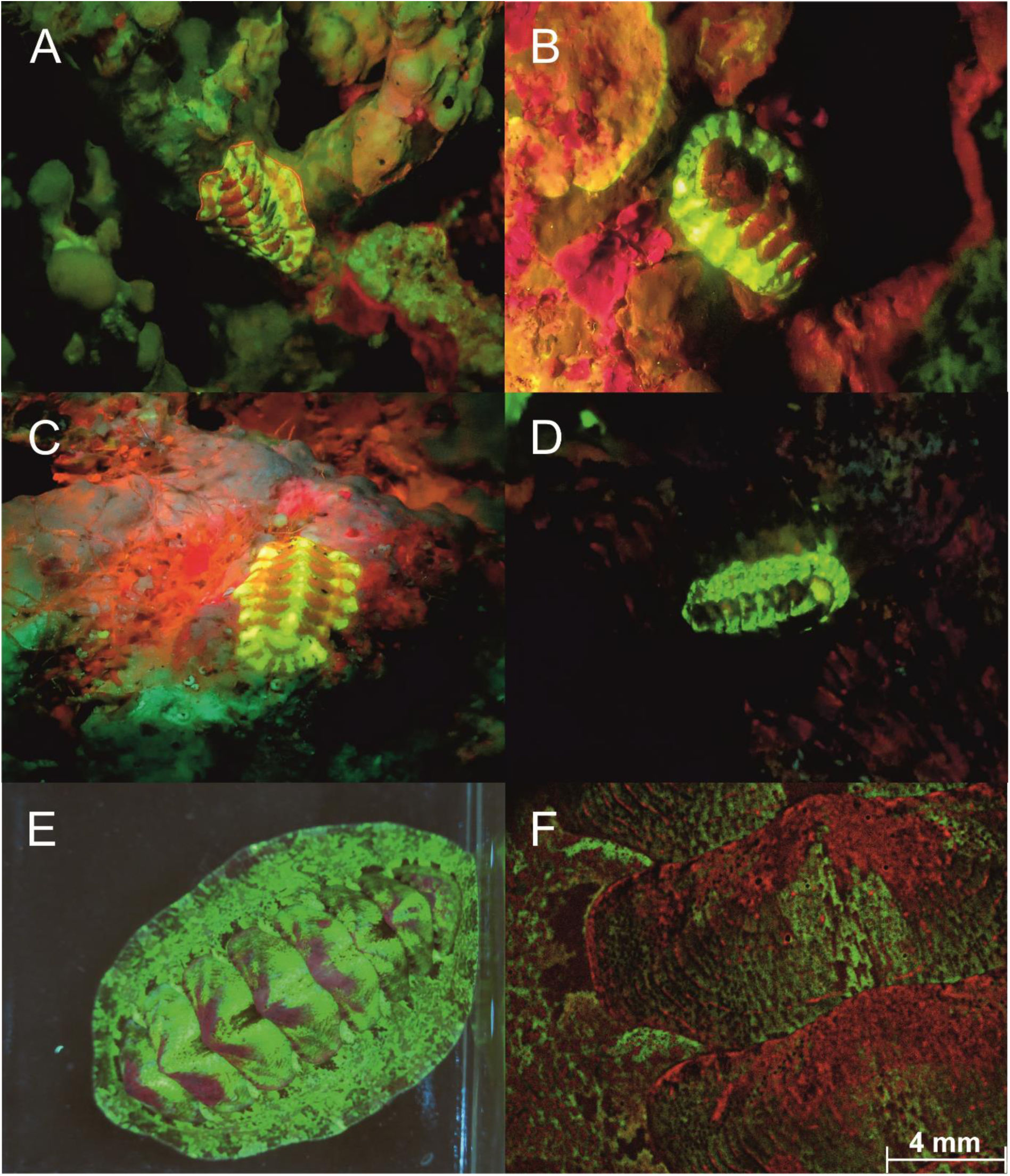
Fluorescence in Polyplacophora: Red, yellow and green fluorescent Polyplacophora in the Red Sea (A-C) and in the Banda Sea (D). Bright green and red fluorescent Polyplacophora in a reef tank in E-F.

Echinodermata – We found bright green, yellow, and red fluorescent crinoid species of different genera in the Banda Sea and in the Red Sea (Figure 7).

**Figure 7:**
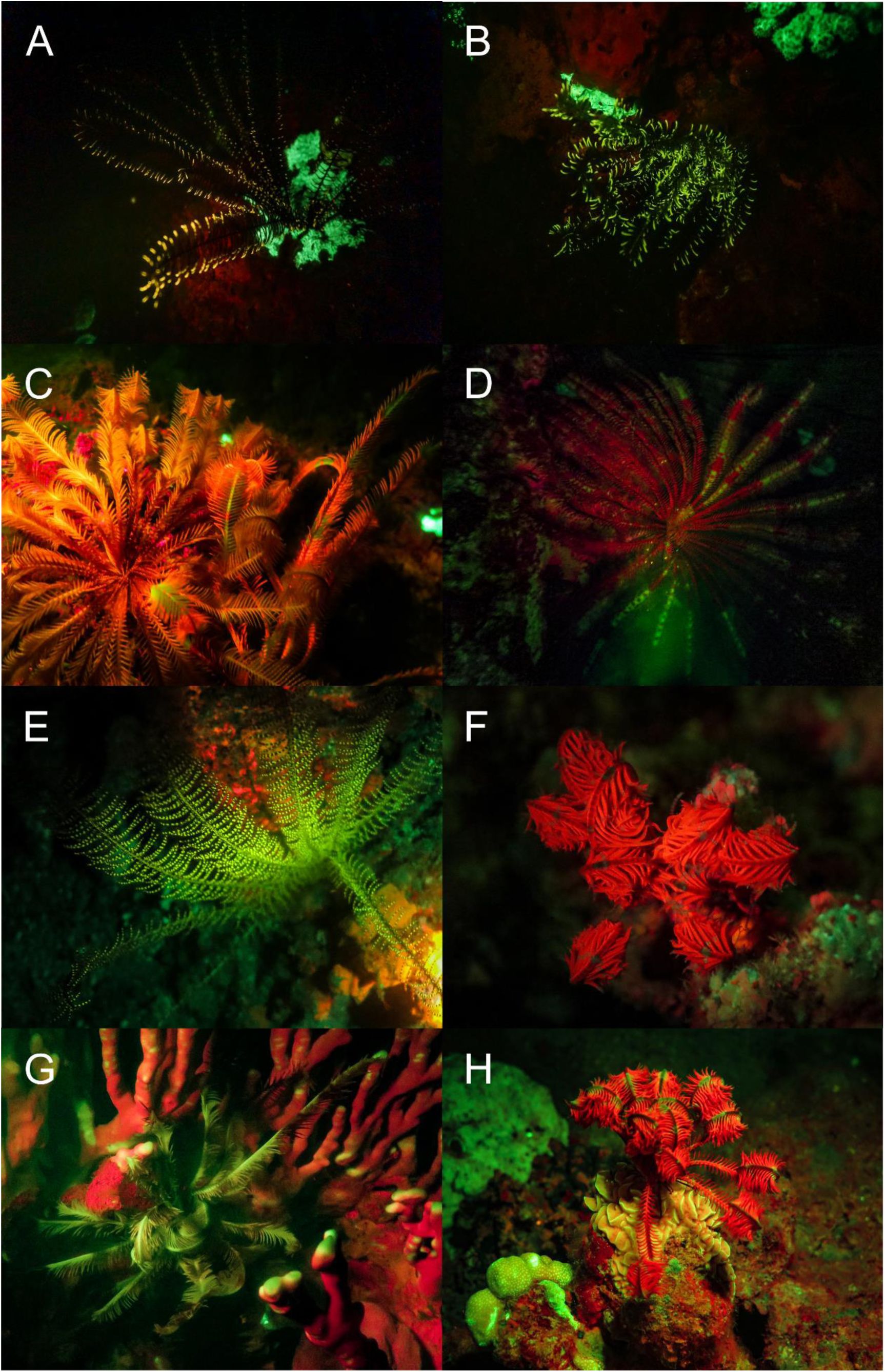
Fluorescence in Echinoderms: New observations of red, yellow and green fluorescence in crinoids from the Banda Sea (A-E) and in the Red Sea (F-H).

Arthropoda - Several fluorescent Decapoda of the family Inachidae, Scyllaridae and, Palaemonidae were found. The decorator crab *Camposcia retusa* (Fig. 8 A found in the Banda Sea showed a bright green fluorescent carapace. In the Red Sea we discovered a bright green fluorescing slipper lobster *Scyllarides sp*. (Fig. 8 B). In addition, we found bright green fluorescence in the carapace of Coleman’s Shrimp*, Periclimenes colemani* (Fig. 8 C), living in symbiosis with fire urchins that do not fluoresce. Green and red fluorescence of the carapace was discovered in peacock mantis shrimps *Odontodactylus scyllarus*. The THUNDER images revealed fluorescence in filamentous structures on maxillipeds (Fig. 9 A, B, F), pleopods (Fig. 9 C), uropods (Fig. 9 E), and on paraeopods (Fig. 9 D) of *Odontodactylus scyllarus*. Furthermore, we identified fluorescent body parts of the mosaic boxer crab (*Lybia tessellata*) (Fig. 9 G).

**Figure 8:**
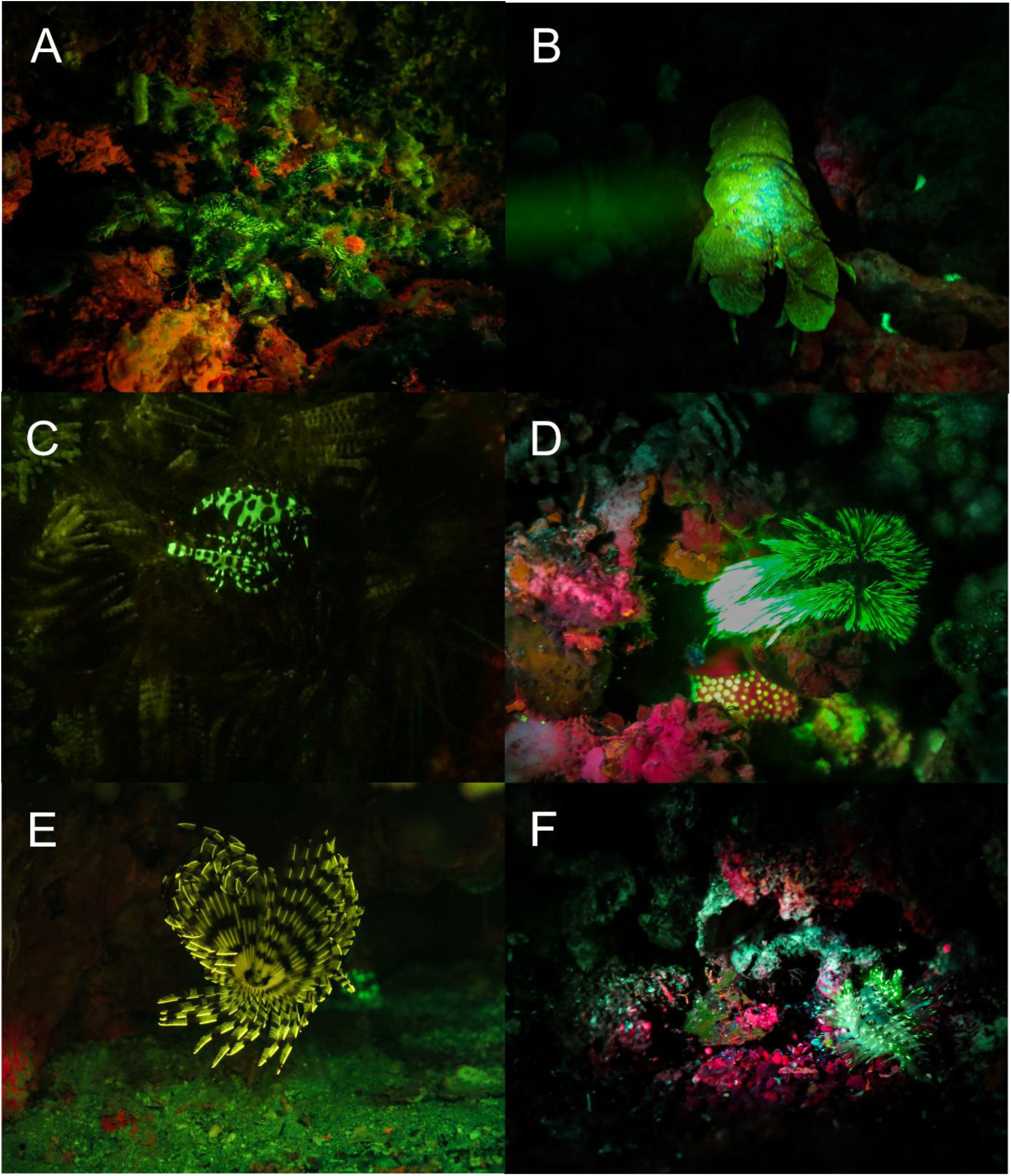
Fluorescence in Arthropoda and Annelida: Green fluorescent *Camposcia retusa* (A)*, Scyllarides sp*. *(B)*, *Periclimenes colemani* (C), Eunicida (D,F), Yellow fluorescent chaeta of an unidentified species of Sabellidae (E). A, C, D, F: Banda Sea, B & E: Red Sea

**Figure 9:**
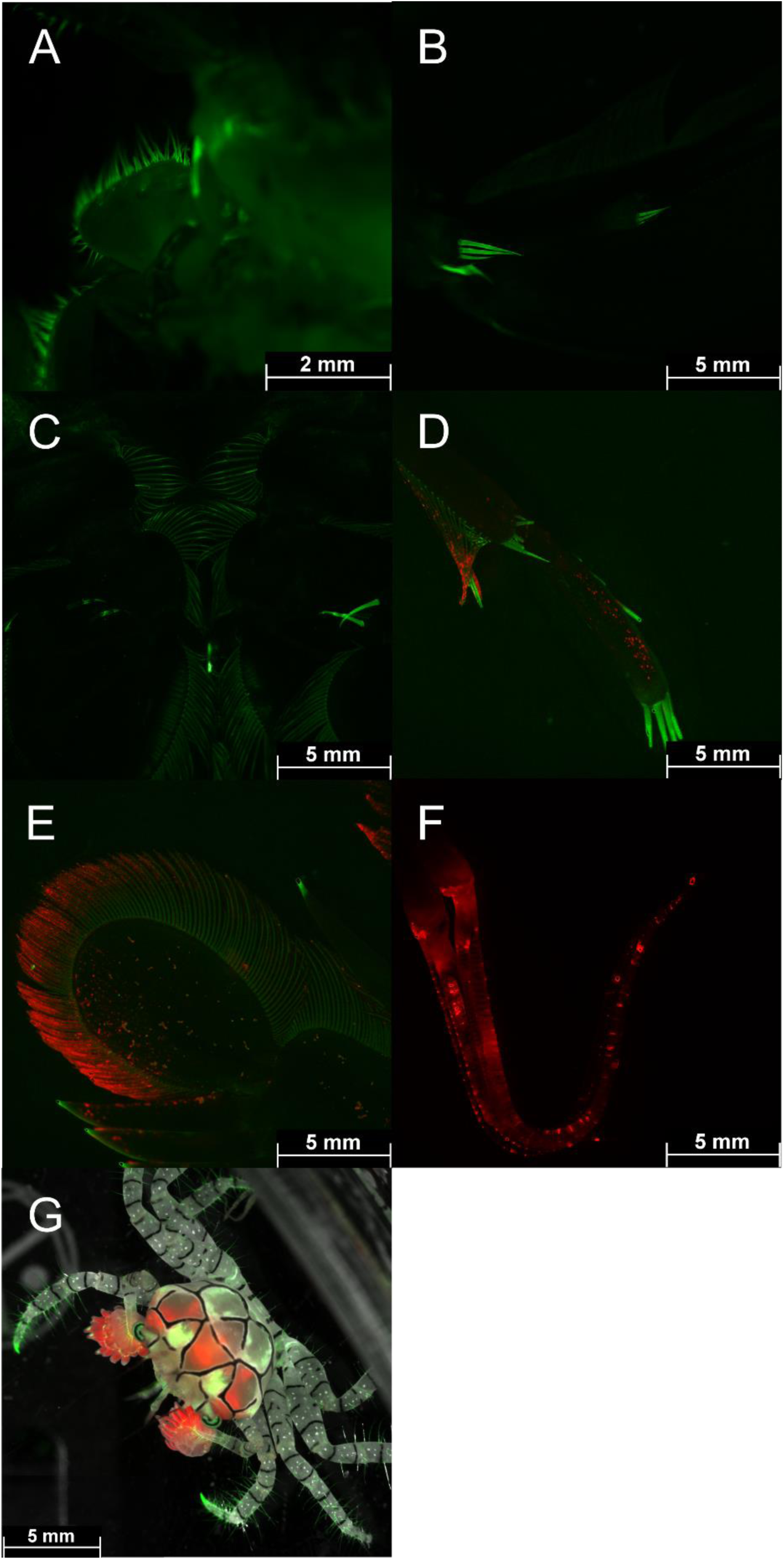
Fluorescence of *Odontodactylus scyllarus* and *Lybia tessalata:* Green (GFP filter) and red fluorescing (Cy5 filter) structures of maxillipeds (A, B, F), pleopods(C), uropods (E), and paraeopods (D). Green and red fluorescing *Lybia tessellata* (G; overlay of brightfield, GFP & CY5 filter). Pictures were taken with a THUNDER microscope (Leica, Germany).

Annelida - We also found fluorescence in two types of bristle worms (polychaetes) from the families Eunicidae and Sabellidae. In the subclass Eunicida only the bristles fluoresced bright green (Fig. 8 D, F). In addition, we found bright yellow fluorescence in a feather duster worm (Sabellidae) (Fig. 8 E).

Chordata – Two fluorescent ascidian species and three species of fluorescent bony fish were photographed in the Banda Sea. Both ascidians *Clavelina coerulea* (Fig. 10 A & B) and *Clavelina robusta* (Fig. 10 C & D) revealed a bright green rim-like fluorescence around their siphons. In both species the fluorescent rims appeared yellow under white light conditions. We found two green and orange fluorescent scarlet frogfish *Antennatus coccineus* (Fig. 11 A & B). The green fluorescence was distributed over the entire body, while the orange fluorescence was found in patches and in the lures. In addition, we found a bright fluorescing green leopard flounder *Bothus pantherinus* (Fig. 11 C). Green fluorescence was more prominent in the white banded sections of the skin. We also found individual differences in fluorescence in the banded pipefish *Corythoichthys intestinalis.* Individuals revealed either green or orange fluorescent skin patterns (Fig. 11 D-F). However, all individuals showed a bright yellow/orange fluorescence around the eyes and the caudal fin (Fig. 11 D-F). Moreover, we observed two individuals of *Pleurosicya mossambica* in the Banda Sea with bright orange-red fluorescence around the eyes and the chorda (Fig. 12 A & B) and two individuals of *Scorpaenopsis possi* with orange and red fluorescent patches in their tissue (Fig. 12 C & D). In addition, we found two different color morphs of *Soleichthys heterorhinos* from the Banda Sea (E) and Red Sea (F) and a green fluorescent undefined species of Lutjanidae (G), and *Brachysomophis henshawi* with a red fluorescent head (H) (Fig. 13 & 14).

**Figure 10:**
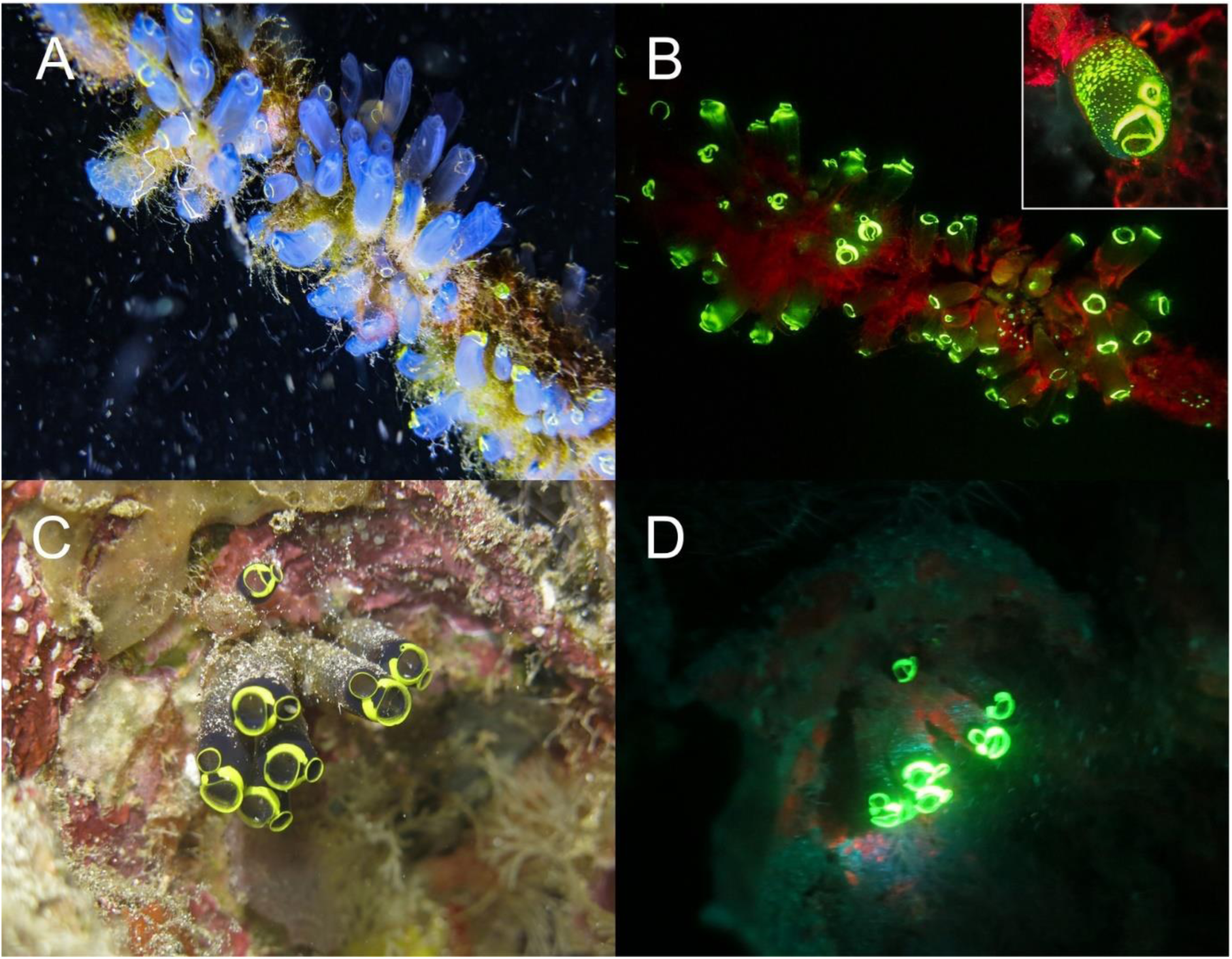
Fluorescence in ascidians in the Banda Sea: White light and fluorescent picture of *Clavelina coerulea* ((A, B) white light in A, fluorescence in B), *Clavelina robusta* ((C, D) white light in C, fluorescence in D). The green fluorescence accumulates on the siphons of the ascidians.

**Figure 11:**
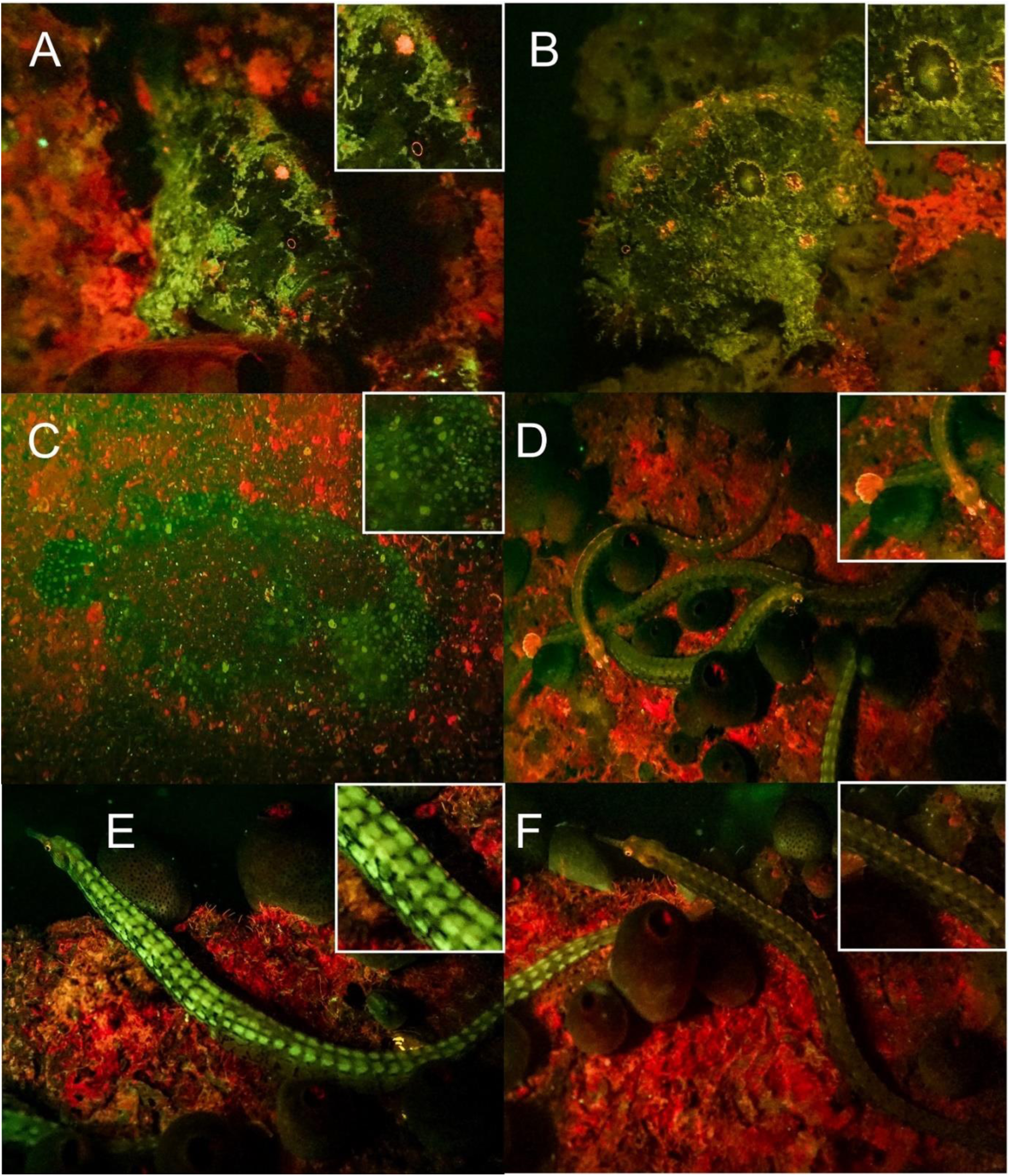
Fluorescence in fish in the Banda Sea: Green and orange fluorescing *Antennatus coccineus* (A, B), green fluorescing *Bothus pantherinus* (C) and different fluorescing individuals of *Corythoichthys intestinalis* (D-F).

**Figure 12:**
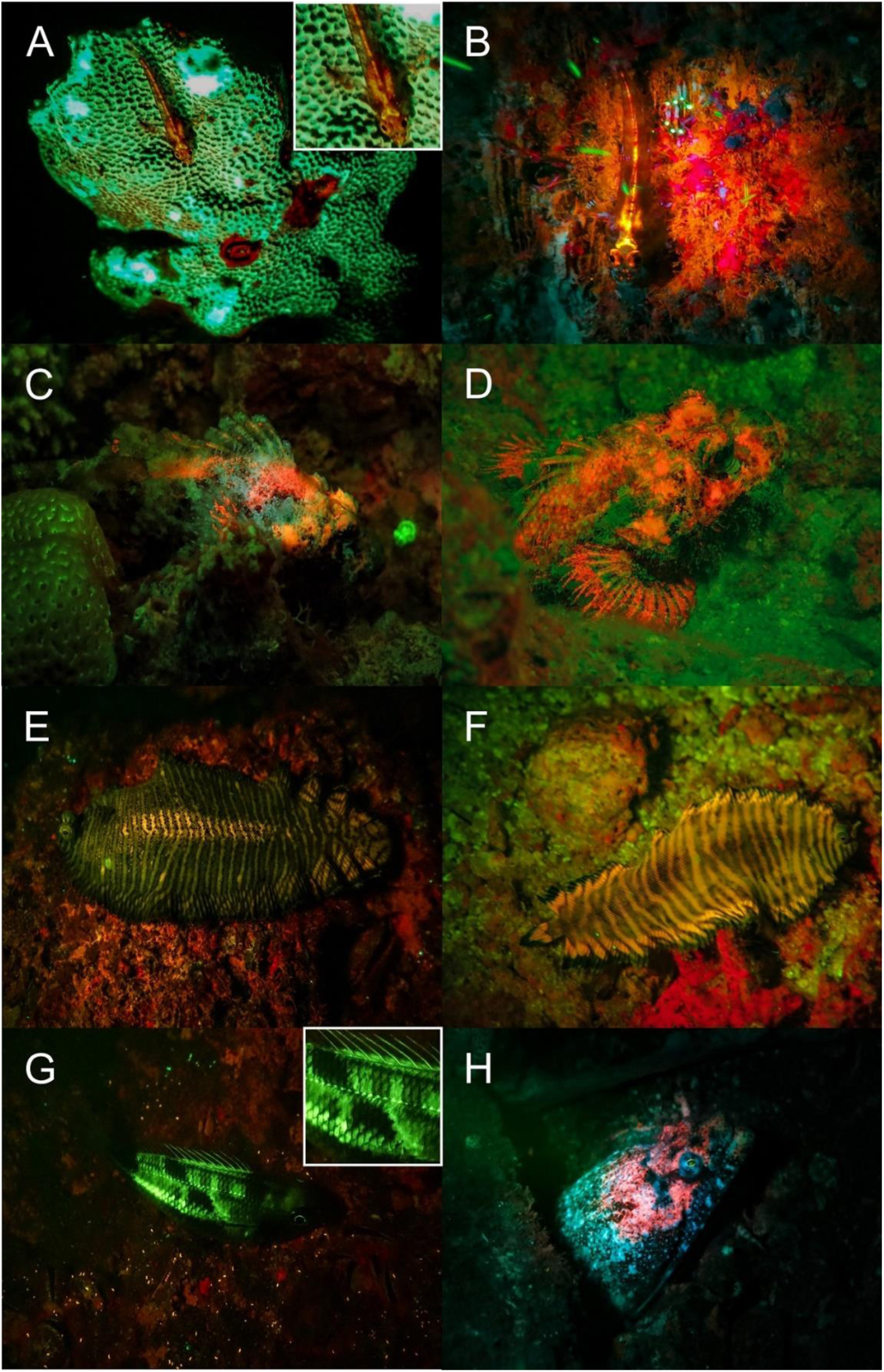
Fluorescence in fish: Red fluorescing *Pleurosicya mossambica* (A & B)*, Scorpaenopsis possi* (C & D), different colormorphs of orange fluorescing *Soleichthys heterorhinos* (E & F), undefined species of Lutjanidae (G), and *Brachysomophis henshawi* (H) with a red fluorescing head. A, B, D, F-H: Banda Sea, C & E: Red Sea.

**Figure 13:**
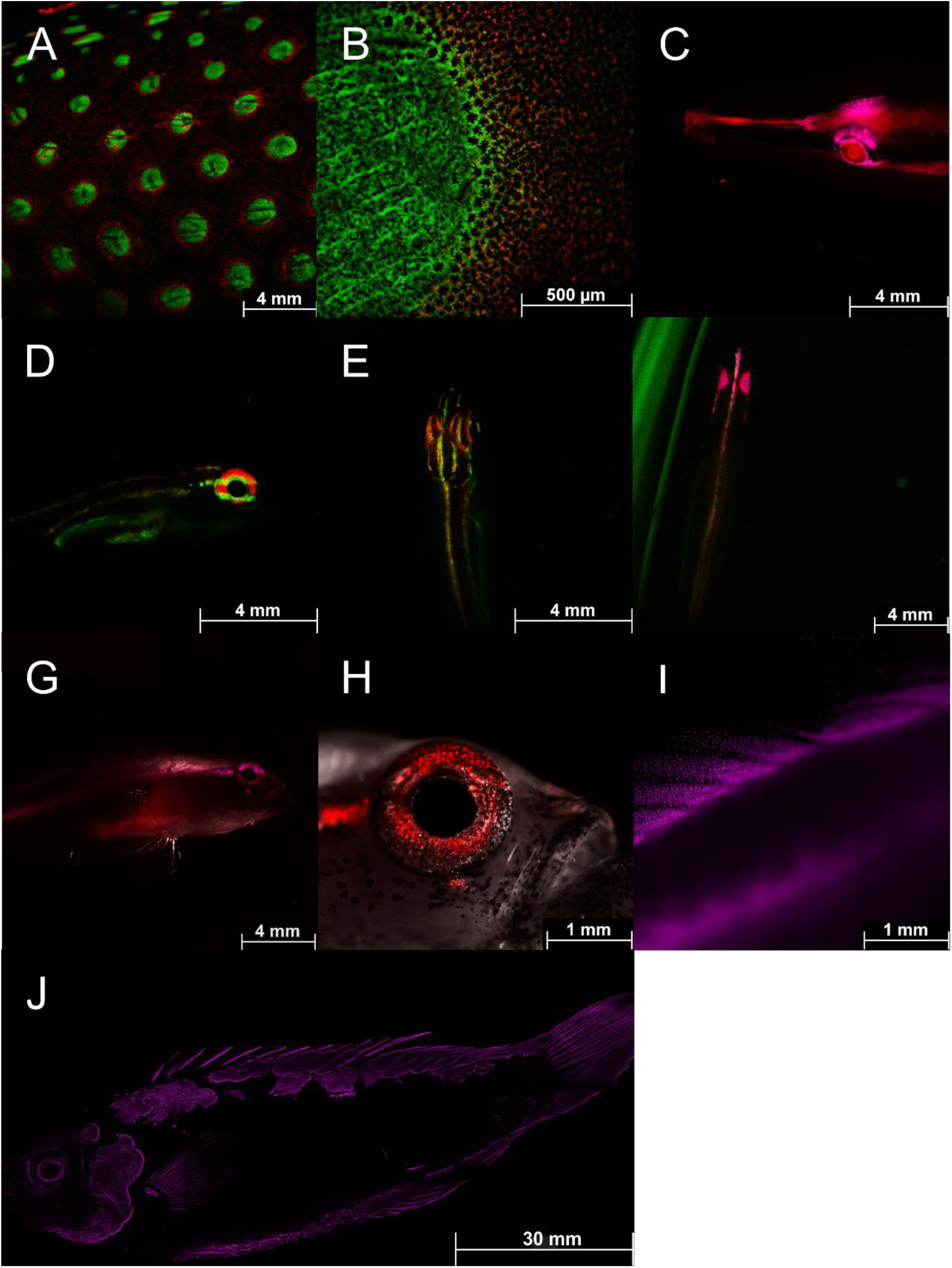
Fluorescence in fish: Detailed fluorescent pictures of *Anampses meleagrides* (A, B; both overlay of GFP and CY5 filter), *Doryrhamphus excisus* (C; overlay of mCherry and CY5 filter), *Eviota atriventris* (D, E; both overlay of GFP & CY5 filter), *Eviota nigriventris* (F-I; F: overlay of GFP, CY5, mCherry filter; G: overlay of mCherry and CY5 filter; H: overlay of brightfield & CY5 filter; I: mCherry filter), *Cirrhilabrus aquamarinus* (J; mCherry filter), Pictures were taken with a Thunder microscope (Leica, Germany).

**Figure 14:**
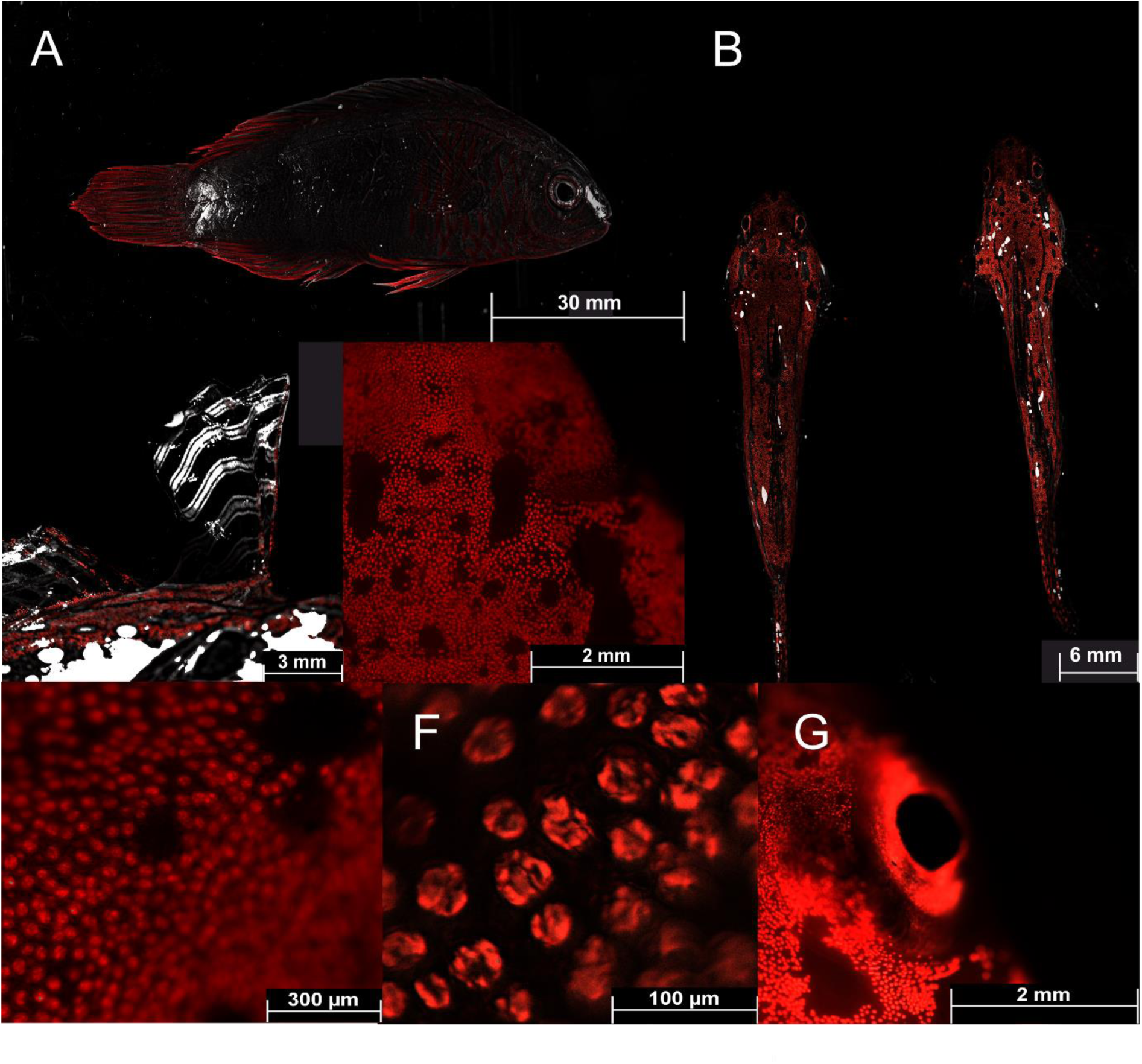
Fluorescence in fish: Detailed fluorescent pictures of *Cirrhilabrus aquamarinus* (A; brightfield & Cy5 filter), Pair of *Synchiropus sycorax* (B; ♀ left and ♂ right; overlay of brightfield and Cy5 filter), first dorsal fin of *Synchiropus sycorax* ♂ (C; overlay of brightfield and Cy5 filter), detailed fluorophores of *Synchiropus sycorax* (D-F; Cy5 filter), fluorescent patch around the eye of *Synchiropus sycorax* ♂ (G; Cy5 filter). Pictures were taken with a Thunder microscope (Leica, Germany).

## Discussion

During night dives in the Banda Sea, Indonesia and the Red Sea, Egypt we photographed marine species, in which, to our knowledge, fluorescence had not been documented in scientific journals before and for which fluorescence has not been characterized. These included species of sponges, crustaceans, polychaetes, slugs, snails, octopus, ascidians, and fish.

Various molecules and anatomical structures that mediate fluorescence have been identified in different species. These include fluorescent proteins, carotenoids, pteridine, porphyrins, tryptophane derivatives, and chlorophyll [2,21,32,33]. Fluorescent proteins have been found in a few taxa of metazoans including Cnidaria and Arthropoda, but also in Cephalochordata and Vertebrata [2,9]. Since we found fluorescence in two species of ascidians that are closer related to vertebrates than other invertebrates, next generation sequencing techniques can reveal if these ascidians contain FP-like genes. In both species bright green fluorescence was seen in a rim-like structure around the oral and atrial siphon (Fig. 6). Since food is ingested via the oral siphon, fluorescence might be involved in attraction of plankton which has been demonstrated for the green fluorescence of the jellyfish *Olindias formosa* [34] and for green and orange fluorescence in corals [35].

In Arthropoda different molecules have been described to be fluorescent [8,36–38]. The blue fluorescence in the carapace of the lobster *Homarus gammarus* is mediated by a multimolecular carotenoprotein, α crustacyanin, binding to the carotenoid astaxanthin [39]. The cuticle of scorpions contains the alkaloid β-carboline which is derived from tryptamine and is excited by UV light leading to the emission of blue fluorescent light [40]. The *Drosophila melanogaster* mutant ‘sepia’ is characterized by red-colored eyes, which is mediated by the accumulation of sepiapterin, a yellow fluorescent pteridine derivative [41,42]. Sepiapterin is involved in the tetrahydrobiopterin pathway essential for the breakdown of phenylalanine, suggesting that this molecule is conserved throughout evolution. The regulation of sepiapterin reductase activity could therefore be a mechanism to accumulate sepiapterin to make specific body parts fluorescent. However, this must be demonstrated. Whether α crustacyanin [39], β-carboline or sepiapterin also contribute to the green fluorescent cuticula of the slipper lobster (Fig. 8 B) and other fluorescent Arthropoda has to be investigated. Because the fluorescence in crustacea and some other species is not intense and only becomes visible with intense excitation, it is difficult to conclude a functional purpose (e.g. Fig 6 C; Fig 8 B, C & F; Fig 9 A, D, E).

Fluorescence has also been described in segmented worms. The intertidal worm *Eulalia sp.* (Polychaeta) secretes a blue-green fluorescent mucus [20]. The gossamer worm (*Tomopteri*s *spp., a* pelagic annelid) uses fluorescence to enhance bioluminescent light [43]. The mucus of the tubeworm *Chaetopterus variopedatus* contains blue and green fluorescence mediated by riboflavin and related derivatives [44,45]. In addition, blue fluorescence and bioluminescence has also been observed in the fireworm *Odontosyllis phosphorea* [46,47]. We also found fluorescence in fire or bristle worms from the families Eunicidae and Sabellidae with so far unknown composition and function (Fig. 5 D-F).

Bioluminescence in some crinoids species is known [48], as well as fluorescent substances derived from crinoids [49]. however, no in situ recordings of fluorescent Crinoidea is known yet. We have found several species in which the fluorescence ranges from yellow/greenish tips of the arms to green/red fluorescent bodies (Fig. 7 A-H). The origin of fluorescence and its possible ecological benefit still need to be investigated.

Various species from different phyla particularly Cnidaria, but also Porifera, Mollusca or Arthropoda live in symbiosis with photosynthetic algae and/or bacteria, which reveal chlorophyll mediated red fluorescence [50–52]. For example, green and orange fluorescence is found in the skeletal elements of a few sponge species. The fluorescence originates most likely from associated algae within the sponge skeletal elements [53]. We also found green, yellow and orange fluorescence in sponges, where the origin and function has not been characterized. Isopods and some molluscs, such as giant clams, also live in symbiosis with red fluorescent, photosynthetic microbes [21].

The fluorescence seen for *Tridacna* is therefore most likely associated to the chlorophyll photosystem II of the algae [50, 52] or involves porphyrins, which are responsible for the red fluorescence in the shells of some gastropd species. In snails, fluorescence is often found in the cerata, which originate either from ingested food (green fluorescence) or symbiotic algae (red fluorescence) [19]. It could be possible, that the nudibranch *Facelina rhodopos* deposits ingested fluorescent food in its cerata since this snail feeds on the hydrozoan *Millepora* that has a weak fluorescence. Behavioural experiments with *Goniobranchus splendidus* revealed that predators learn to avoid feeding on the unpalatable snail. The avoidance behaviour of the fish is triggered by the yellow rim of the snail [54]. The red fluorescent rim of *Nembrotha kubaryana* may, therefore, have similar functions. While ingested or attached symbionts explain the red fluorescence of snails and giant clams, the chemical composition of yellow and red fluorescent rims in *Goniobranchus splendidus* and *Nembrotha kubaryana* has to be investigated.

Green and red fluorescence is also common in fish [55]. Red fluorescence around 600 nm, for example, has been reported in over 30 reef fish from more than 16 genera and 5 families [56]. The red fluorescence has been associated with guanine crystals, frequently produced in iridophores and has been found in pipefish (Fig.11), triplefins, blennies, and gobies. Red fluorescence is found in the iris and parts of the head and thorax and more rarely in fins. It has been shown that the black-faced blenny *Tripterygion delaisi,* which has red fluorescent irises can perceive and respond to red fluorescence [13–15]. Perception of possible fluorescent rivals leads to an increase in aggressive behavior in the red-eyed wrasse *Cirrhilabrus solorensis* [11,12,16]. In crypto-benthic fish, such as the scarlet frogfish (*Antennatus coccineus*) and the leopard flounder (*Bothus pantherinus*) (Figure 11) fluorescence may facilitate background matching (camouflage) [7,55]. This has recently been demonstrated in scorpionfish for adjusting red fluorescence to background luminance [57].

We also photographed yellow fluorescent corals that live near the water surface (≤ 15 m (Fig. 3). So far, no yellow GFP-like fluorescent protein has been isolated and cloned from stony corals [5]. Also, it was rather known that yellow fluorescent corals can occur in mesophotic reefs [58]. This could be useful for further studies on fluorescent proteins from stony corals. While the isolation and characterization of new yellow fluorescent proteins would be desirable, it has to be mentioned that the coexpression and colocalization of green and red fluorescent proteins may also lead to yellow fluorescence [59].

In summary, this study describes fluorescent marine organisms of different species in which fluorescence has not been published before in scientific literature. A total of 27 species, in which fluorescence has not been described in scientific literature before are added to the list of known fluorescent marine organisms. Three of these species belong to the phylum Porifera, seven to the phylum Mollusca, three to the subphylum Crustacea, three to the phylum of Annelida, two to the class Ascidiacea, and three to the subphylum Vertebrata. We describe the first cases of fluorescence in Octopoda and Ascidiacea, and also show fluorescence in Nudibranchia, where the fluorescence – unlike most previous observations - is not linked to ingested food. This study, therefore, extends the palette of fluorescence in marine species. It shows that fluorescence likely is a common phenomenon and that its diversity is not limited to cnidarians. Systematically searching marine biodiversity hotspots with blue or UV lights will, therefore, likely result in the discovery of more fluorescent species and molecules, which will help understand the diverse roles fluorescence may play in marine ecosystems.

## Acknowledgements

We thank all RUB students who contributed to this article by taking underwater pictures. In particular, we thank Lukas Menzen, Andre Marcel Haase, Leonard Klaus, Denise Wagner, and Jan-Niklas Schnalke. Thanks to the dive instructors and dive guides who showed us fluorescence in coral reefs during night dives.

## Availability of data and material

All data generated or analysed during this study are included in this published article [and its supplementary information files].

## Authors’ contributions

Conceptualization LHP, MH, SH

Investigation LHP, BS, GL

Writing – Original LHP, BS, GL, MH, SH

Writing – Review & Editing LHP, PJ, MH, SH

## Supplementary material 1

Example for digital processing of the images and details on camera specifications and settings.

## Supplementary material 2

Table of 27 fluorescent species we described in this paper.

## References

1. Johnsen S. The Optics of Life. A Biologist’s Guide to Light in Nature. Princeton: Princeton University Press; 2011.

2. Macel M-L, Ristoratore F, Locascio A, Spagnuolo A, Sordino P, D’Aniello S. Sea as a color palette: the ecology and evolution of fluorescence. Zoological Lett. 2020; 6:9. Epub 2020/06/10. doi: 10.1186/s40851-020-00161-9 PMID: 32537244.

3. Ferreira G, Bollati E, Kühl M. The role of host pigments in coral photobiology. Front Mar Sci. 2023; 10. doi: 10.3389/fmars.2023.1204843.

4. Shimomura O. Discovery of Green Fluorescent Protein. In: Chalfie M, editor. Green fluorescent protein. Properties, applications and protocols. 2nd ed. Hoboken, NJ: Wiley-Interscience; 2006. pp. 1–13.

5. Alieva NO, Konzen KA, Field SF, Meleshkevitch EA, Hunt ME, Beltran-Ramirez V, et al. Diversity and evolution of coral fluorescent proteins. PLOS ONE. 2008; 3:e2680. Epub 2008/07/16. doi: 10.1371/journal.pone.0002680 PMID: 18648549.

6. Chudakov DM, Matz MV, Lukyanov S, Lukyanov KA. Fluorescent proteins and their applications in imaging living cells and tissues. Physiological Reviews. 2010; 90:1103–63. doi: 10.1152/physrev.00038.2009 PMID: 20664080.

7. Sparks JS, Schelly RC, Smith WL, Davis MP, Tchernov D, Pieribone VA, et al. The covert world of fish biofluorescence: a phylogenetically widespread and phenotypically variable phenomenon. PLoS One. 2014; 9:e83259. Epub 2014/01/08. doi: 10.1371/journal.pone.0083259 PMID: 24421880.

8. Shagin DA, Barsova EV, Yanushevich YG, Fradkov AF, Lukyanov KA, Labas YA, et al. GFP-like proteins as ubiquitous metazoan superfamily: evolution of functional features and structural complexity. Mol Biol Evol. 2004; 21:841–50. Epub 2004/02/12. doi: 10.1093/molbev/msh079 PMID: 14963095.

9. Kumagai A, Ando R, Miyatake H, Greimel P, Kobayashi T, Hirabayashi Y, et al. A bilirubin-inducible fluorescent protein from eel muscle. Cell. 2013; 153:1602–11. Epub 2013/06/13. doi: 10.1016/j.cell.2013.05.038 PMID: 23768684.

10. Bomati EK, Manning G, Deheyn DD. Amphioxus encodes the largest known family of green fluorescent proteins, which have diversified into distinct functional classes. BMC Evol Biol. 2009; 9:77. Epub 2009/04/21. doi: 10.1186/1471-2148-9-77 PMID: 19379521.

11. Gerlach T, Sprenger D, Michiels NK. Fairy wrasses perceive and respond to their deep red fluorescent coloration. Proc Biol Sci. 2014; 281. doi: 10.1098/rspb.2014.0787 PMID: 24870049.

12. Gerlach T, Theobald J, Hart NS, Collin SP, Michiels NK. Fluorescence characterisation and visual ecology of pseudocheilinid wrasses. Front Zool. 2016; 13:13. Epub 2016/03/15. doi: 10.1186/s12983-016-0145-1 PMID: 26981144.

13. Wucherer MF, Michiels NK. Regulation of red fluorescent light emission in a cryptic marine fish. Front Zool. 2014; 11:1. Epub 2014/01/08. doi: 10.1186/1742-9994-11-1 PMID: 24401080.

14. Kalb N, Schneider RF, Sprenger D, Michiels NK. The Red-Fluorescing Marine Fish *Tripterygion delaisi* can Perceive its Own Red Fluorescent Colour. Ethology. 2015; 121:566–76. doi: 10.1111/eth.12367.

15. Bitton P-P, Harant UK, Fritsch R, Champ CM, Temple SE, Michiels NK. Red fluorescence of the triplefin *Tripterygion delaisi* is increasingly visible against background light with increasing depth. R Soc Open Sci. 2017; 4:161009. Epub 2017/03/22. doi: 10.1098/rsos.161009 PMID: 28405391.

16. Anthes N, Theobald J, Gerlach T, Meadows MG, Michiels NK. Diversity and Ecological Correlates of Red Fluorescence in Marine Fishes. Front Ecol Evol. 2016; 4. doi: 10.3389/fevo.2016.00126.

17. Liaci L. Natura e localizzazione di un pigmento fluorescents in *Aaptos aaptos* O. S. (Demospongiae). Bolletino di zoologia. 1962; 29:425–31. doi: 10.1080/11250006209440630.

18. Read KR, Davidson JM, Twarog BM. Fluorescence of sponges and coelenterates in blue light. Comparative Biochemistry and Physiology. 1968; 25:873–82. doi: 10.1016/0010-406X(68)90575-6.

19. Betti F, Bavestrello G, Cattaneo-Vietti R. Preliminary evidence of fluorescence in Mediterranean heterobranchs. Journal of Molluscan Studies. 2021; 87. doi: 10.1093/mollus/eyaa040.

20. Rodrigo AP, Lopes A, Pereira R, Anjo SI, Manadas B, Grosso AR, et al. Endogenous Fluorescent Proteins in the Mucus of an Intertidal Polychaeta: Clues for Biotechnology. Mar Drugs. 2022; 20. Epub 2022/03/25. doi: 10.3390/md20040224 PMID: 35447897.

21. Lindquist N, Barber PH, Weisz JB. Episymbiotic microbes as food and defence for marine isopods: unique symbioses in a hostile environment. Proc Biol Sci. 2005; 272:1209–16. doi: 10.1098/rspb.2005.3082 PMID: 16024384.

22. Burgess WE, Axelrod HR, Hunziker RE. Dr. Burgess’s Atlas of marine aquarium fishes. Neptune City, NJ: T.F.H. Publ; 1990.

23. Debelius H. Fischführer Indischer Ozean. Rotes Meer bis Thailand ; über 900 Farbfotos aus dem natürlichen Lebensraum der Meeresfische. 2nd ed. Melle: Tetra; 1996.

24. Patzner RA, Baensch HA. Perciformes (Barschartige). Ausgenommen Falter- und Kaiserfische. 1st ed. Melle: Mergus Verl. für Natur- und Heimtierkunde Baensch; 1998.

25. Patzner RA, Mossleitner H. Non-Perciformes (Nicht-Barschartige). Sowie Falter- und Kaiserfische. 1st ed. ; 1999.

26. Erhardt H. Wirbellose. 1st ed. Melle: Mergus Verlag für Natur- und Heimtierkunde Baensch; 1998.

27. Erhardt H. Wirbellose. 1st ed. Melle: Mergus Verlag für Natur- und Heimtierkunde Baensch; 2000.

28. Debelius H. Fisch-Führer Indopazifik. Malediven bis Philippinen ; 700 Fischarten in ihrem natürlichen Lebensraum fotografiert. 1st ed. Münster: NTV; 2002.

29. Debelius H, Kuiter RH. Nacktschnecken der Weltmeere. 1200 Arten weltweit. 1st ed. Stuttgart: Kosmos; 2007.

30. Kuiter RH, Debelius H. Atlas der Meeresfische. Die Fische an den Küsten der Weltmeere. 3rd ed. Stuttgart: Kosmos; 2009.

31. Kuiter RH, Debelius H. Atlas der wirbellosen Meerestiere. Weichtiere, Würmer, Stachelhäuter, Krebstiere. Stuttgart: Kosmos; 2009.

32. Gaevskii NA, Kolmakov VI, Dubovskaya OP, Klimova EP. Interrelations of Epibiontic Microalgae and Crustacean Zooplankton under Conditions of a Blooming Eutrophic Water Body. Russian Journal of Ecology. 2004; 35:35–41. doi: 10.1023/B:RUSE.0000011107.72097.1c.

33. Wicksten MK. A Review and a Model of Decorating Behavior in Spider Crabs (Decapoda, Brachyura, Majidae). Crustac. 1993; 64:314–25. doi: 10.1163/156854093x00667.

34. Haddock SHD, Dunn CW. Fluorescent proteins function as a prey attractant: experimental evidence from the hydromedusa *Olindias formosus* and other marine organisms. Biol Open. 2015; 4:1094–104. Epub 2015/07/31. doi: 10.1242/bio.012138 PMID: 26231627.

35. Ben-Zvi O, Lindemann Y, Eyal G, Loya Y. Coral fluorescence: a prey-lure in deep habitats. Commun Biol. 2022; 5:537. Epub 2022/06/02. doi: 10.1038/s42003-022-03460-3 PMID: 35654953.

36. Masuda H, Takenaka Y, Yamaguchi A, Nishikawa S, Mizuno H. A novel yellowish-green fluorescent protein from the marine copepod, *Chiridius poppei*, and its use as a reporter protein in HeLa cells. Gene. 2006; 372:18–25. Epub 2006/02/14. doi: 10.1016/j.gene.2005.11.031 PMID: 16481130.

37. Evdokimov AG, Pokross ME, Egorov NS, Zaraisky AG, Yampolsky IV, Merzlyak EM, et al. Structural basis for the fast maturation of Arthropoda green fluorescent protein. EMBO Rep. 2006; 7:1006–12. Epub 2006/08/25. doi: 10.1038/sj.embor.7400787 PMID: 16936637.

38. Hunt ME, Scherrer MP, Ferrari FD, Matz MV. Very bright green fluorescent proteins from the Pontellid copepod *Pontella mimocerami*. PLOS ONE. 2010; 5:e11517. Epub 2010/07/14. doi: 10.1371/journal.pone.0011517 PMID: 20644720.

39. Chayen NE, Cianci M, Grossmann JG, Habash J, Helliwell JR, Nneji GA, et al. Unravelling the structural chemistry of the colouration mechanism in lobster shell. Acta Crystallogr D Biol Crystallogr. 2003; 59:2072–82. Epub 2003/11/27. doi: 10.1107/s0907444903025952 PMID: 14646064.

40. Stachel SJ, Stockwell SA, van Vranken DL. The fluorescence of scorpions and cataractogenesis. Chemistry & Biology. 1999; 6:531–9. doi: 10.1016/s1074-5521(99)80085-4 PMID: 10421760.

41. Forrest H, Nawa S. Structures of Sepiapterin and Isosepiapterin. Nature. 1962; 196:372–3. doi: 10.1038/196372b0.

42. Kim J, Suh H, Kim S, Kim K, Ahn C, Yim J. Identification and characteristics of the structural gene for the Drosophila eye colour mutant sepia, encoding PDA synthase, a member of the omega class glutathione S-transferases. Biochem J. 2006; 398:451–60. doi: 10.1042/BJ20060424 PMID: 16712527.

43. Gouveneaux A, Flood PR, Erichsen ES, Olsson C, Lindström J, Mallefet J. Morphology and fluorescence of the parapodial light glands in *Tomopteris helgolandica* and allies (Phyllodocida: Tomopteridae). Zoologischer Anzeiger. 2017; 268:112–25. doi: 10.1016/j.jcz.2016.08.002.

44. Deheyn DD, Enzor LA, Dubowitz A, Urbach JS, Blair D. Optical and physicochemical characterization of the luminous mucous secreted by the marine worm *Chaetopterus sp*. Physiol Biochem Zool. 2013; 86:702–5. Epub 2013/10/16. doi: 10.1086/673869 PMID: 24241067.

45. Branchini BR, Behney CE, Southworth TL, Rawat R, Deheyn DD. Chemical Analysis of the Luminous Slime Secreted by the Marine Worm *Chaetopterus* (Annelida, Polychaeta). Photochem Photobiol. 2014; 90:247–51. Epub 2013/10/17. doi: 10.1111/php.12169 PMID: 24004150.

46. Deheyn DD, Latz MI. Internal and secreted bioluminescence of the marine polychaete *Odontosyllis phosphorea* (Syllidae). Invertebrate Biology. 2009; 128:31–45. doi: 10.1111/j.1744-7410.2008.00149.x.

47. Meulenaere E de, Puzzanghera C, Deheyn DD. Self-powered bioluminescence in a marine tube worm. The FASEB Journal. 2020; 34:1. doi: 10.1096/fasebj.2020.34.s1.04618.

48. Mallefet J, Martinez-Soares P, Eléaume M, O’Hara T, Duchatelet L. New insights on crinoid (Echinodermata; Crinoidea) bioluminescence. Front Mar Sci. 2023; 10. doi: 10.3389/fmars.2023.1136138.

49. Singh AJ, Gorka AP, Bokesch HR, Wamiru A, O’Keefe BR, Schnermann MJ, et al. Harnessing Natural Product Diversity for Fluorophore Discovery: Naturally Occurring Fluorescent Hydroxyanthraquinones from the Marine Crinoid *Pterometra venusta*. J Nat Prod. 2018; 81:2750–5. Epub 2018/11/29. doi: 10.1021/acs.jnatprod.8b00761 PMID: 30495954.

50. Pearse VB, Muscatine L. Role of symbiotic algae (Zooxanthellae) in coral calcification. The Biological Bulletin. 1971; 141:350–63. doi: 10.2307/1540123.

51. Trench RK, Wethey DS, Porter JW. Observations on the symbiosis with zooxanthaellae among the Tridacnidae (Molussca, Bivalvia). The Biological Bulletin. 1981; 161:180–98. doi: 10.2307/1541117.

52. Trautman DA, Hinde R. Sponge/Algal Symbioses: A Diversity of Associations. In: Seckbach J, editor. Symbiosis. Mechanisms and Model Systems. Secaucus: Kluwer Academic Publishers; 2002. pp. 521–37.

53. Steyaert M, Mogg A, Dunn N, Dowell R, Head CEI. Observations of coral and cryptobenthic sponge fluorescence and recruitment on autonomous reef monitoring structures (ARMS). Coral Reefs. 2022; 41:877–83. doi: 10.1007/s00338-022-02283-2.

54. Winters AE, Green NF, Wilson NG, How MJ, Garson MJ, Marshall NJ, et al. Stabilizing selection on individual pattern elements of aposematic signals. Proc Biol Sci. 2017; 284. doi: 10.1098/rspb.2017.0926 PMID: 28835556.

55. Brauwer M de, Hobbs J-PA, Ambo-Rappe R, Jompa J, Harvey ES, McIlwain JL. Biofluorescence as a survey tool for cryptic marine species. Conserv Biol. 2018; 32:706–15. Epub 2018/02/19. doi: 10.1111/cobi.13033 PMID: 28984998.

56. Michiels NK, Anthes N, Hart NS, Herler J, Meixner AJ, Schleifenbaum F, et al. Red fluorescence in reef fish: a novel signalling mechanism. BMC Ecol. 2008; 8:16. Epub 2008/09/16. doi: 10.1186/1472-6785-8-16 PMID: 18796150.

57. John L, Santon M, Michiels NK. Scorpionfish rapidly change colour in response to their background. Front Zool. 2023; 20:10. Epub 2023/03/03. doi: 10.1186/s12983-023-00488-x PMID: 36864453.

58. Eyal G, Wiedenmann J, Grinblat M, D’Angelo C, Kramarsky-Winter E, Treibitz T, et al. Spectral Diversity and Regulation of Coral Fluorescence in a Mesophotic Reef Habitat in the Red Sea. PLOS ONE. 2015; 10:e0128697. Epub 2015/06/24. doi: 10.1371/journal.pone.0128697 PMID: 26107282.

59. Koyama-Honda I, Ritchie K, Fujiwara T, Iino R, Murakoshi H, Kasai RS, et al. Fluorescence imaging for monitoring the colocalization of two single molecules in living cells. Biophys J. 2005; 88:2126–36. Epub 2004/12/13. doi: 10.1529/biophysj.104.048967 PMID: 15596511.

