## Supplemental Table 1 for "New observations of fluorescent organisms in the Banda Sea and in the Red Sea"

**Supporting information_1**

Example for the digital processing

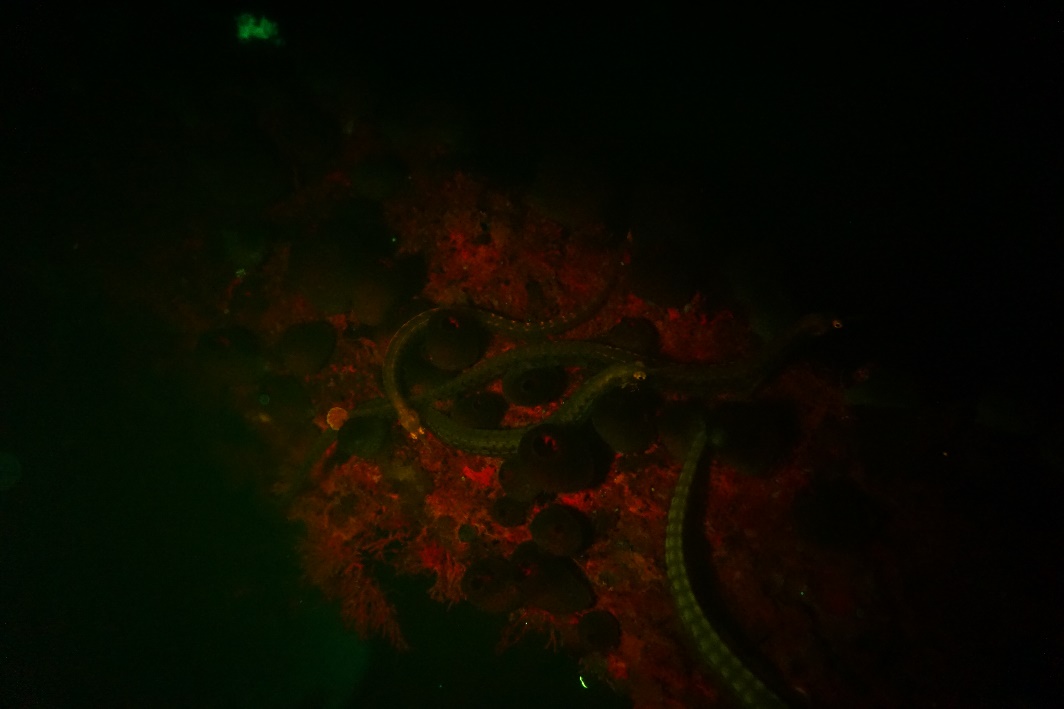

Figure 1: Raw-data image. Sony α 6000, F/3,5, 1/80 sec, ISO: 3200, WB: Underwater-Auto

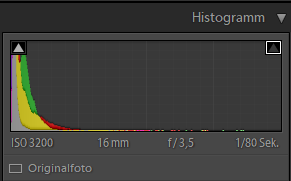

Figure 2: Histogram of the Raw-data picture in Lightroom.

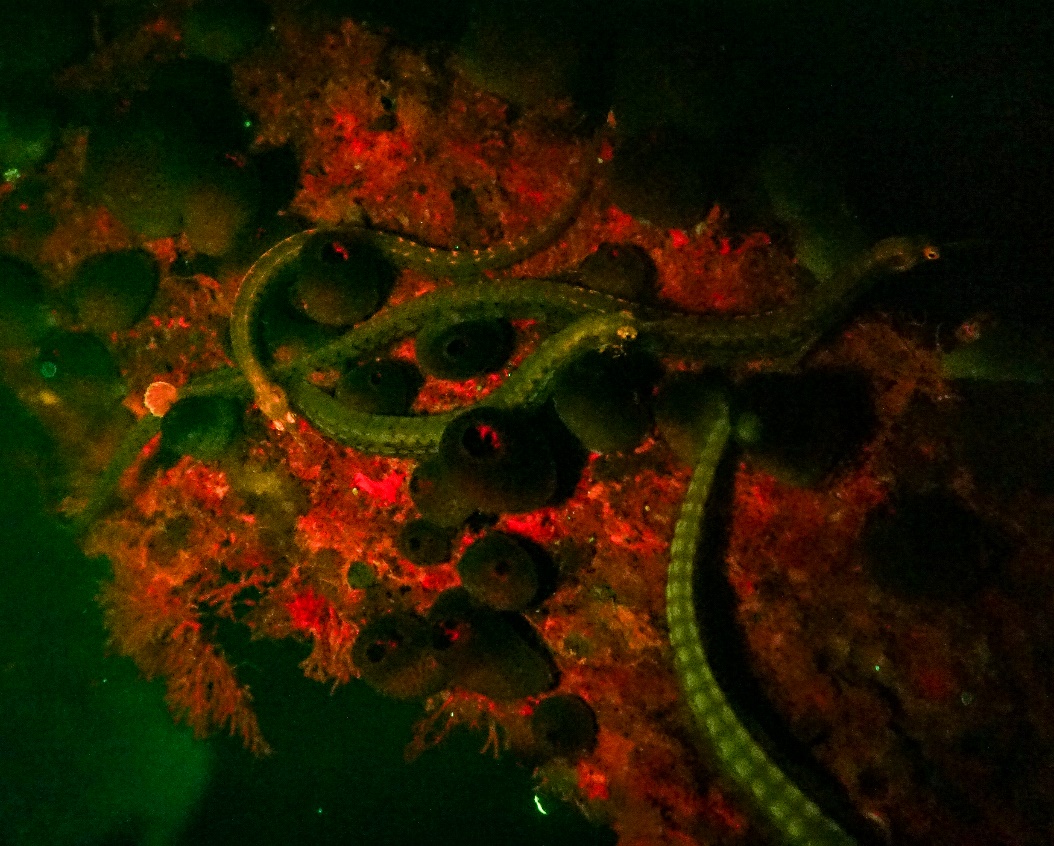

Figure 3: Edited image: cropping and increasing contrast (+69) and exposure (+2,09)

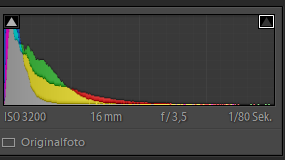

Figure 4: Histogram of the edited image in Lightroom

Underwater-image details on camera specifications and settings:

Figure 2:

A: Canon Poweshot G15, F/2.5, 1/160 sec., ISO 80, WB automatic;

B: Canon Powershot G15, F/2, 1/25 sec., ISO 800, WB underwater automatic; light source: Sola light source

C: Canon Powershor G 15, F/2, 1/500 sec., ISO 12800, WB underwater automatic; light source: Sola light

D: Sony α 6000, F/3.5, 1/125 sec., ISO 3200, WB underwater automatic; light source: Sola light

Figure 3:

A: Canon Powershot G15, F/1.8, 1/30 sec., ISO 320, WB underwater automatic, light source: Sola light

B: Canon Powershot G15, F/1.8, 1/30 sec., ISO 400, WB underwater automatic, light source: Sola light

C: Canon Powershot G15, F/2.8, 1/60 sec., ISO 640, WB underwater automatic; light source: Ex-Inon Flashlight

D: Canon Powershot G15, F/2.5, 1/25 sec., ISO 800, WB underwater automatic, light source: Sola light

E: Canon Powershot G15, F/3.5, 1/160 sec., ISO 500, WB underwater automatic, light source: Sola light

F: Canon Powershot G15, F/1.8, 1/20 sec., ISO 1250, WB underwater automatic; light source: Ex-Inon Flashlight

Figure 4:

A: Canon Powershot G15, F/1.8, 1/100 sec., ISO 400, WB automatic

B: Screenshot of Sony α 6000 Video, WB underwater automatic; light source: Sola light

C: Canon Powershot G15, F/1.8, 1/15 sec., ISO 1600, WB automatic

D: Sony α 6000, F/3.5, 1/20 sec., ISO 3200, WB underwater automatic; light source: Sola light

E: Canon Powershot G15, F/2.2, 1/50 sec., ISO 1600, WB underwater automatic; light source: Sola light

F: Sony α 6000, F/3.5, 1/40 sec., ISO 3200, WB underwater automatic; light source: Sola light

G: Canon Powershot G15, F/2.8, 1/30 sec., ISO 800, WB underwater automatic; light source: Sola light

H: Canon Powershot G15, F/2.5, 1/60 sec., ISO 640, WB underwater automatic; light source: Ex-Inon Flashlight

Figure 5:

A: Canon Powershot G15, F/2.2, 1/60 sec., ISO 160, WB underwater automatic

B: Sony α 6000, F/7.1, 1/60 sec., ISO 1600, WB automatic; light source: Sola light

Figure 6:

A: Canon Powershot G15, F/2.8, 1/60 sec., ISO 640, WB underwater automatic; light source: Ex-Inon Flashlight

B: Canon Powershot G15, F/6.5, 1/80 sec., ISO 3200, WB underwater automatic; light source: Sola light

C: Canon Powershot G15, F/2.5, 1/100 sec., ISO 500, WB underwater automatic; light source: Blue Star light

D: Canon Powershot G15, F/2.8, 1/8 sec., ISO 1600, WB underwater automatic; light source: Blue Star

Figure 7:

A: Sony α 6000, F/3.5, 1/80 sec., ISO 3200, WB underwater automatic; light source: Blue Star light

B: Sony α 6000, F/3.5, 1/160 sec., ISO 3200, WB underwater automatic; light source: Sola light

C: Canon Powershot G15, F/1.8, 1/20 sec., ISO 1250, WB underwater automatic; light source: Sola light

D: Canon Powershot G15, F/1.8, 1/8 sec., ISO 1600, WB underwater automatic; light source: Sola light

E: Canon Powershot G15, F/2, 1/8 sec., ISO 1600, WB underwater automatic; light source: Sola light

F: Canon Powershot G15, F/2.8, 1/160 sec., ISO 1600, WB underwater automatic; light source: Sola light

G: Canon Powershot G15, F/3.5, 1/25 sec., ISO 3200, WB underwater automatic; light source: Sola light

H: Canon Powershot G15, F/2.2, 1/60 sec., ISO 640, WB underwater automatic; light source: Ex-Inon Flashlight

Figure 8:

A: Canon Powershot G15, F/2.2, 1/8 sec., ISO 1600, WB underwater automatic; light source: Sola light

B: Canon Powershot G15, F/2.2, 1/10 sec., ISO 1600, WB underwater automatic; light source: Blue Star light

C: Sony α 6000, F/3.5, 1/100 sec., ISO 3200, WB underwater automatic; light source: Sola light

D: Canon Powershot G15, F/2.2, 1/40 sec., ISO 800, WB underwater automatic; light source: Sola light

E: Canon Powershot G15, F/2.8, 1/60 sec., ISO 640, WB underwater automatic; light source: Sola light

F: Canon Powershot G15, F/2.2, 1/20 sec., ISO 1000, WB underwater automatic; light source: Sola light

Figure 10:

A: Canon Powershot G15, F/1.8, 1/30 sec., ISO 160, WB automatic

B: Sony α 6000, F/3.5, 1/100 sec., ISO 3200, WB underwater automatic; light source: Sola light

C: Canon Powershot G15, F/2.8, 1/200 sec., ISO 320, WB automatic

D: Canon Powershot G15, F/2, 1/20 sec., ISO 800, WB underwater automatic; light source: Blue star light

Figure 11:

A: Sony α 6000, F/3.5, 1/50 sec., ISO 3200, WB underwater automatic; light source: Sola light

B: Sony α 6000, F/3.5, 1/40 sec., ISO 3200, WB underwater automatic; light source: Sola light

C: Screenshot of Sony α 6000 Video, WB underwater automatic; light source: Sola light

D: Sony α 6000, F/3.5, 1/80 sec., ISO 3200, WB underwater automatic; light source: Sola light

E: Sony α 6000, F/3.5, 1/80 sec., ISO 3200, WB underwater automatic; light source: Sola light

F: Sony α 6000, F/3.5, 1/80 sec., ISO 3200, WB underwater automatic; light source: Sola light

Figure 12:

A: Sony α 6000, F/4, 1/60 sec., ISO 6400, WB underwater automatic; light source: Sola light

B: Canon Powershot G15, F/1.8, 1/30 sec., ISO 500, WB underwater automatic; light source: Sola light

C: Canon Powershot G15, F/2.2, 1/160 sec., ISO 1600, WB underwater automatic; light source: Ex-Inon Flashlight

D: Canon Powershot G15, F/2.8, 1/60 sec., ISO 640, WB underwater automatic; light source: Ex-Inon Flashlight

E: Screenshot of Sony α 6000 Video, WB underwater automatic; light source: Sola light

F: Canon Powershot G15, F/2.5, 1/15 sec., ISO 1600, WB underwater automatic; light source: Sola light

G: Sony α 6000, F/3.5, 1/80 sec., ISO 3200, WB underwater automatic; light source: Sola light

H: Canon Powershot G15, F/2.2, 1/20 sec., ISO 1600, WB underwater automatic; light source: Blue star light
