## Supplemental Table 2 for "New observations of fluorescent organisms in the Banda Sea and in the Red Sea"

**Supporting information_2**

Table 1: Fluorescent species described in the paper, in which fluorescence has not been described yet in peer-reviewed literature.

| **Fluorescent species** | **Figure** |
| --- | --- |
| *Gelliodes fibulata* | 2 A & B |
| unidentified sponge | 2 C |
| unidentified sponge | 2 D |
| *Abdopus aculeatus* | 4 A & B |
| *Euprotomus bulla* | 4 F |
| *Facelina rhodopus* | 4 H |
| *Nebrotha kubaryana* | 5 |
| unidentified Polyplacophora | 6 A - C |
| unidentified Polyplacophora | 6 D |
| unidentified Polyplacophora | 6 E & F |
| Unidentified Crinoid | 7 A & B |
| Unidentified Crinoid | 7 C & D |
| Unidentified Crinoid | 7 E |
| Unidentified Crinoid | 7 F & H |
| Unidentified Crinoid | 7 G |
| *Camposcia retusa* | 8 A |
| *Scyllarides sp*. | 8 B |
| *Periclimenes colemani* | 8 C |
| Unidentified Sabellidae | 8 E |
| *Odontodactylus scyllarus* | 9 A -F |
| *Lybia tessalata* | 9 G |
| *Clavelina coerulea* | 10 A & B |
| *Clavelina robusta* | 10 C & D |
| *Antennatus coccineus* | 11 A & B |
| *Bothus pantherinus* | 11 C |
| *Scorpaenopsis possi* | 12 C & D |
| *Brachysomophis henshawi* | 12 H |
